# Widespread ancient whole genome duplications in Malpighiales coincide with Eocene global climatic upheaval

**DOI:** 10.1101/215608

**Authors:** Liming Cai, Zhenxiang Xi, André M. Amorim, M. Sugumaran, Joshua S. Rest, Liang Liu, Charles C. Davis

**Affiliations:** Department of Organismic and Evolutionary Biology, Harvard University Herbaria, 22 Divinity Avenue, Cambridge, Massachusetts 02138, USA; Key Laboratory of Bio-resource and Eco-environment of Ministry of Education, College of Life Sciences, Sichuan University, Chengdu 610064, China; Departamento de Ciências Biológicas, Universidade Estadual de Santa Cruz, Ilhéus, 45.662-900, Bahia, Brazil; Rimba Ilmu Botanic Garden, Institute of Biological Sciences, University of Malaya, 50603 Kuala Lumpur, Malaysia; Department of Ecology and Evolution, Stony Brook University, Stony Brook, New York, USA; Department of Statistics and Institute of Bioinformatics, University of Georgia, Athens, GA 30602, USA

**Keywords:** climatic upheaval, flowering plants, genome evolution, global change, phylogenomics, speciation

## Abstract

Ancient whole genome duplications (WGDs) are important in eukaryotic genome evolution, and are especially prominent in plants. Recent genomic studies from large vascular plant clades, including ferns, gymnosperms, and angiosperms suggest that WGDs may represent a crucial mode of speciation. Moreover, numerous WGDs have been dated to events coinciding with major episodes of global and climatic upheaval, including the mass extinction at the KT boundary (~65 Ma) and during more recent intervals of global aridification in the Miocene (~10-5 Ma). These findings have led to the hypothesis that polyploidization may buffer lineages against the negative consequences of such disruptions. Here, we explore WGDs in the large, and diverse flowering plant clade Malpighiales using a combination of transcriptomes and complete genomes from 42 species. We conservatively identify 22 ancient WGDs, widely distributed across Malpighiales subclades. Our results provide strong support for the hypothesis that WGD is an important mode of speciation in plants. Importantly, we also identify that these events are clustered around the Eocene-Paleocene Transition (~54 Ma), during which time the planet was warmer and wetter than any period in the Cenozoic. These results establish that the Eocene Climate Optimum represents another, previously unrecognized, period of prolific WGDs in plants, and lends support to the hypothesis that polyploidization promotes adaptation and enhances plant survival during major episodes of global change. Malpighiales, in particular, may have been particularly influenced by these events given their predominance in the tropics where Eocene warming likely had profound impacts owing to the relatively tight thermal tolerances of tropical organisms.

**Significance Statement:** Whole genome duplications (WGDs) are hypothesized to generate adaptive variations during episodes of climate change and global upheaval. Using large-scale phylogenomic assessments, we identify an impressive 22 ancient WGDs in the large, tropical flowering plant clade Malpighiales. This supports growing evidence that ancient WGDs are far more common than has been thought. Additionally, we identify that WGDs are clustered during a narrow window of time, ~54 Ma, when the climate was warmer and more humid than during any period in the last ~65 Ma. This lends support to the hypothesis that WGDs are associated with surviving climatic upheavals, especially for tropical organisms like Malpighiales, which have tight thermal tolerances.

---

Whole genome duplication (WGD), or polyploidy, is an important evolutionary force that has shaped plant evolution. It has long been appreciated that the formation of recent polyploids in vascular plants is common (1, 2), and mounting evidence suggests that ancient polyploids are more frequent than once thought. Well-cited examples of ancient WGDs have been associated with the origin of several hyperdiverse clades, including in the common ancestor of seed plants, flowering plants, monocots, orchids, core eudicots, mustards (Brassicaceae), legumes (Fabaceae), and sunflowers (Asteraceae) (3-13). In addition, WGDs have been identified in ferns and gymnosperms (14, 15), thus expanding the phylogenetic scope of this phenomenon to span vascular plants. Among these ancient WGDs, several have been dated to the Cretaceous-Tertiary (KT) boundary (~65 Ma), potentially linking these polyploidization events to plants’ abilities to survive abrupt global environmental change (16, 17). Similarly, a large number of WGDs have also been reported in grasses during the late Miocene when arid, grass dominated landscapes expanded dramatically (18). In these cases, the potential adaptive value of WGDs is thought to arise from the origin of genetic novelties (19-22) and by masking the effects of deleterious mutations (23), which may facilitate plant survival across periods of global disruption. Although debate exists as to the influence of WGD on speciation and enhanced species diversification rates (14, 24-28), it is generally accepted that chromosomal rearrangements from WGDs can significantly accelerate isolating barriers, thus promoting cladogenesis (29-31). In short, it is established that WGDs are a prominent feature of vascular plant evolution, but the respective phylogenetic distribution, timing, and significance of these ancient events remains unclear.

Here, we investigate WGDs in the large and diverse angiosperm order Malpighiales, which contains more than 16,000, mostly tropical, species with tremendous morphological and ecological diversity. Members of this clade also include numerous economically important crops such as rubber, cassava, and flax. The Malpighiales have long been recognized as one of the most difficult clades to resolve in the flowering plant tree of life (32, 33), which has been attributed in part to its rapid radiation in the mid-Cretaceous (33, 34). However, recent efforts utilizing phylogenomic approaches have greatly increased our understanding of deep level relationships within the order (34). Additionally, the clade includes numerous species that have previously been targeted for genomic investigation of WGDs. Eight genomes are currently available for interrogation, including *Hevea brasiliensis* (rubber), *Manihot esculenta* (cassava), *Linum usitatissimum* (flax), *Ricinus communis* (castor bean)*, Jatropha curcas* (Barbados nut), *Populus trichocarpa* (black cottonwood), *Salix suchowensis* (shrub willow), and *Salix purpurea* (purple willow). Notably, three WGDs have previously been identified in Malpighiales using these data: in the common ancestor of *Populus* and *Salix* (35-65 Ma; 35, 36); in the common ancestor of *Manihot* and *Hevea* (35-47 Ma; 37); and more recently in *Linum* (5-9 Ma; 38). In addition, studies using transcriptomes have identified an older WGD shared by other blue-flowered *Linum* species at 20-40 Ma (33). More complicated polyploidy histories involving multiple rounds of WGDs and hybridization have also been reported using chromosome count data in the genera *Passiflora* (39) and *Viola* (40, 41). In summary, owing to the apparently considerable propensity and frequency of WGDs in Malpighiales, combined with existing complete sequence data available for the order, the clade is an ideal study system for investigating the frequency and timing of WGDs.

## Results and Discussion

We utilized a multi-pronged, phylogenomic pipeline to rigorously identify, locate, and determine the age of WGDs in Malpighiales using three lines of inference that have been commonly applied in angiosperms and elsewhere: rates of synonymous substitutions (Ks) among paralogs (e.g., 4, 42), phylogenomic (gene tree) reconciliation (e.g., 10, 15), and a likelihood based gene count method (e.g., 43, 44). Our data set included 36 ingroup taxa derived from eight genomes and 28 transcriptomes (15 newly acquired for this study) plus six outgroup species (Table S1-3). This sampling includes 21 traditionally recognized families in the order, thus representing the broad outline of phylodiversity within Malpighiales (*sensu* APG IV, 32, 45). An initial Ks analysis was used to identify evidence of WGD for each species. This method identifies a proliferation of duplicated genes from WGDs under the assumption that synonymous substitutions between duplicate genes accrue at a relatively constant rate. Next, we applied a modified version of the phylogenomic pipeline by Jiao et al. (10) to more finely place WGDs among Malpighiales subclades. This method localizes WGDs based on the presence of duplicated genes at particular nodes across a phylogeny. We adapted this method to better accommodate incomplete lineage sorting by incorporating the duplication-transfer-loss model as implemented in *Notung* v2.9 (46). Our final method is a statistically rigorous test of ancient WGDs based on gene count data (43), which estimates the likelihood of putative WGDs on a phylogeny using a gene count matrix summarized across all orthologous genes. This latter approach is advantageous because it suffers less from false positive rates due to tandem duplication and assembly error (43).

*Massive WGDs in Malpighiales–*Our analyses identify an impressive 22–24 WGDs broadly distributed across the Malpighiales phylogeny (Fig. 1). In nearly all cases these events are corroborated by all three methods. Our Ks analysis identified WGD in 22 species (Fig. 1, Fig. S1-2); our phylogenomic reconciliation analysis identified 24 WGDs (Table S4); and 22 WGDs were verified with the likelihood-based gene count method (Table S4). Moreover, these results are robust to data quality and phylogenetic uncertainty (see also Materials and Methods). We additionally analyzed a reduced data set containing only completely sequenced genomes and very high quality transcriptomes to further verify these results. Even with this more conservative data set, we still identified 22 WGDs using all three methods (Table S5). Three of the WGDs we identify validate those from previous studies in *Populus* (35), *Salix* (47, 48), *Manihot* (37), *Hevea* (49), and *Linum* (38). In addition, recent analyses have inferred two independent WGDs in *Linum* using Ks distributions (38, 50). These events, however, have not been corroborated simultaneously due to the limited power of this method. Our reconciliation and gene-count method, in contrast, identifies these two duplications in the lineage leading to *Linum*. Furthermore, we verified these two independent WGDs using a novel, phylogeny-guided synteny analysis (Fig. S3) (9). Here, we identified 15 syntenic regions across large scaffolds reflecting the four-parted paralogous relationship created by two independent WGDs in the *Linum* lineage.

**Figure 1.**
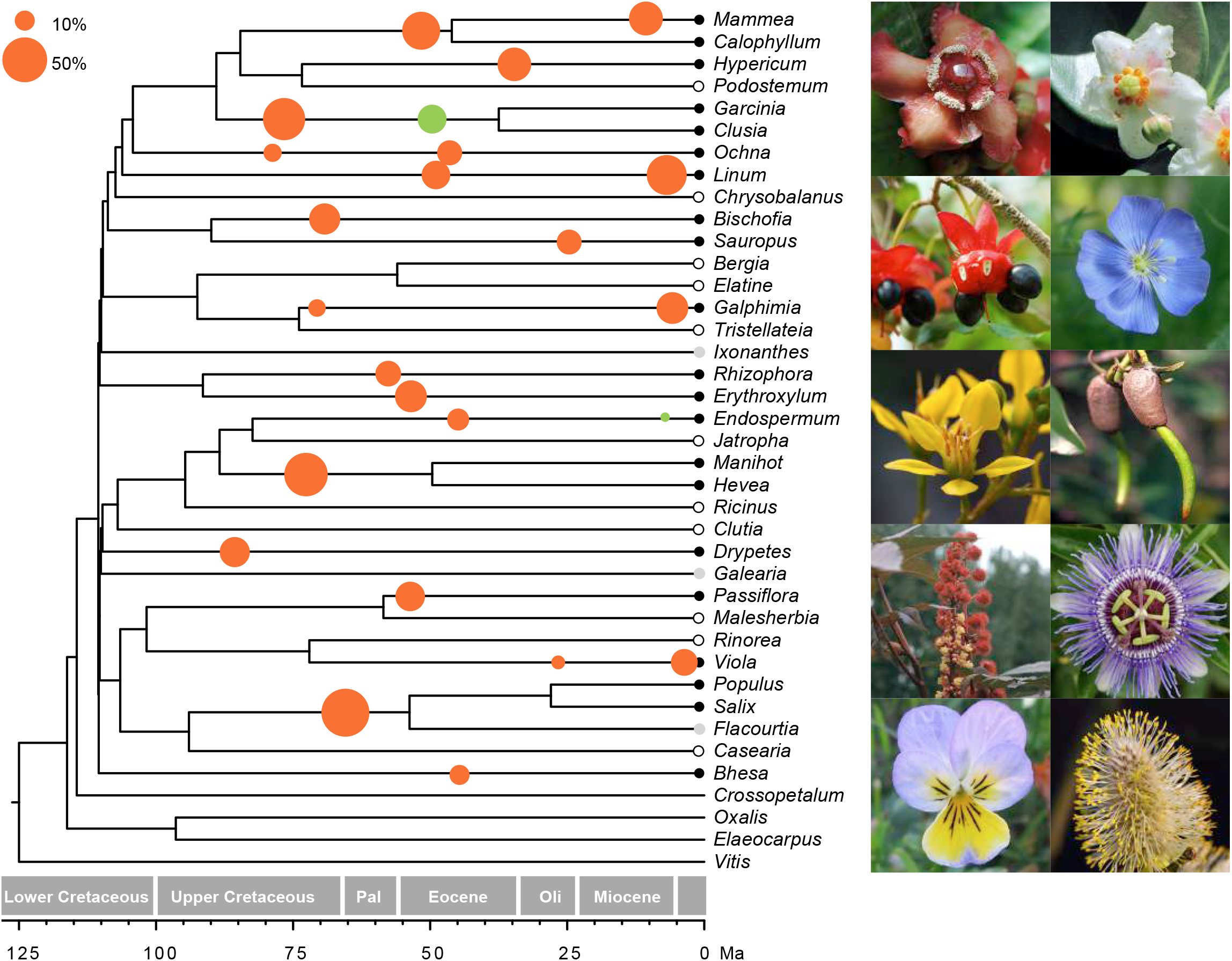
Phylogenetic distribution of WGDs in Malpighiales. Species tree of Malpighiales inferred from 5113 gene trees using a summary coalescent method. WGDs identified from Ks analysis are illustrated with solid black dots on the terminal branches of corresponding species; species indicated with grey dots have transcriptomes that are potentially insufficient for adequate assessment of WGD using the Ks method. Two *Salix* species are collapsed into one terminal branch for simplicity. Circles illustrated along branches are dated WGDs (divergence time of parental genomes) from our phylogenomic reconciliation analysis. The radius of each circle is proportional to the percentage of orthologus genes supporting the WGD as determined from phylogenomic reconciliation (scale, top left). Solid red circles are significant WGDs as confirmed by our gene-count analysis; solid green circles indicate WGDs that do not receive significant support using the gene count method.

One WGD requires more detailed exploration. In our phylogenetic reconciliation and gene-count analysis, a WGD is inferred to predate the origin of the common ancestor of *Populus, Salix*, and *Flacourtia* (the Ks analysis is inconclusive for *Flacourtia*; Fig. 1, Fig. S2). Chromosome count data, however, *do not* support such an early occurrence of WGD. Instead, the chromosome number of *Populus* and *Salix* are approximately twice that of *Flacourtia* (2n = 38; versus 2n =20 or 22 in *Flacourtia*; IPCN Chromosome Reports, http://www.tropicos.org/Project/IPCN) suggesting that this WGD event likely occurred more recently, and is thus restricted to the common ancestor of *Populus* and *Salix*. A similar discrepancy in yeast involving the identification of an older WGD inferred using phylogenetic reconciliation versus a more recent WGD inferred using a gold standard synteny-guided genome comparison (51, 52). Here, the authors provide reasonable evidence that the older WGD is spurious and confounded by an allopolyploidization event resulting from hybridization. The nature of this allopolyploidization resulted in a deeper phylogenetic placement of the WGD. It is important to recognize here that gene-tree data are limited in some respects since they provide an estimate of the divergence times of the parental diploid genomes, but are less conclusive around exactly when the hybridization and polyploidization events occurred (53, 54); dates in Table S4 are thus likely slightly older than the actual polyploidization events. In light of these results, a plausible hypothesis is that the WGD shared by *Populus* and *Salix* results from an allopolyploidization in which an ancestor of the *Flacourtia* lineage served as one parental lineage. Testing and evaluating this hypothesis remains a challenge (55), and is an obvious avenue for future research. Regardless, we do not anticipate this phenomenon to be a pervasive problem for our analysis given that the origin of viable polyploids derived from widely disparate phylogenetic lineages appears to be rare, and thus not likely to greatly influence our placement and dating of the large number of WGDs.

The vast majority of the WGDs we identify have not previously been reported. These WGDs are broadly distributed across 18 branches of the Malpighiales phylogeny. Interestingly, these events are commonly associated with the most diverse clades in the order, including in the clusioids, ochnoids, euphorbioids, phyllanthoids, violets, and passion flowers. Additionally, we note that some species-poor clades show no evidence of duplication (e.g., *Malesherbia*, *Rinorea*, and Elatinaceae, among others; Fig. 1), despite having species-rich sister clades. This lends tentative support to suggestions that WGDs may fuel species diversification (27), possibly via the establishment of reproductive barriers (31). However, other studies have concluded that although polyploidization is important to cladogenesis in plants, it likely does not enhance species diversification rates (14, 18). We cannot fully address this outstanding question with existing data (28), but regardless, our analyses set the stage for establishing the finer scale taxon sampling necessary to pinpoint these events to clarify the association between WGDs and the tempo of diversification in Malpighiales. Namely, do WGDs precede prolific diversification of Malpighiales subclades?

Finally, an obvious question emerges from these results: Are Malpighiales unique among angiosperms in their propensity for pervasive and widespread WGDs? We think that the answer to this question is almost certain to be no. Recent and ongoing investigations incorporating vast nuclear genomic data and extended taxon sampling indicate that other similarly diverse clades, including Asteraceae (4, 11), Poaceae (18), and Caryophyllales (56, 57) also show histories characterized by prolific WGDs. Collectively, these results suggest that Malpighiales and others are likely to represent the tip of the iceberg, thus establishing WGDs as a pervasive phenomenon characteristic of possibly hundreds of major angiosperm clades (and extending to other vascular plant clades, including gymnosperms and ferns—14, 15). Further investigation is required to address this question more broadly, but this seems a plausible hypothesis in light of recent findings.

*Timing of WGDs coincides with events of climate upheaval*–In addition to determining the phylogenetic placement of WGDs in Malpighiales we also estimated the divergence time of WGD-derived paralogs using penalized likelihood. Our results indicate that the timing of WGDs range broadly from 5.7-85.0 Ma (Fig. 2, Table S4). Surprisingly, however, these events are not randomly distributed in time. Instead, our Gaussian mixture model (58) indicates that the inferred ages of WGDs display a striking bimodal distribution (Bayesian Information Criterion, BIC −225.05, compared to BIC −227.30 for the univariate normal distribution) with peaks at the Eocene-Paleocene (mean age 53.89 Ma) and late Miocene (mean age 7.39 Ma), respectively. Our assessment using bootstrap replicates provide confident statistical support for this interpretation: 98% of the replicates support a bimodal distribution of WGDs with the mean age of each cluster ranging from 6.06–13.78 Ma and 52.26–57.79 Ma. In our case, an overwhelming number of WGDs occur during the older, early Eocene time period, where there are 19 events, versus only five during the more recent late Miocene period.

**Figure 2.**
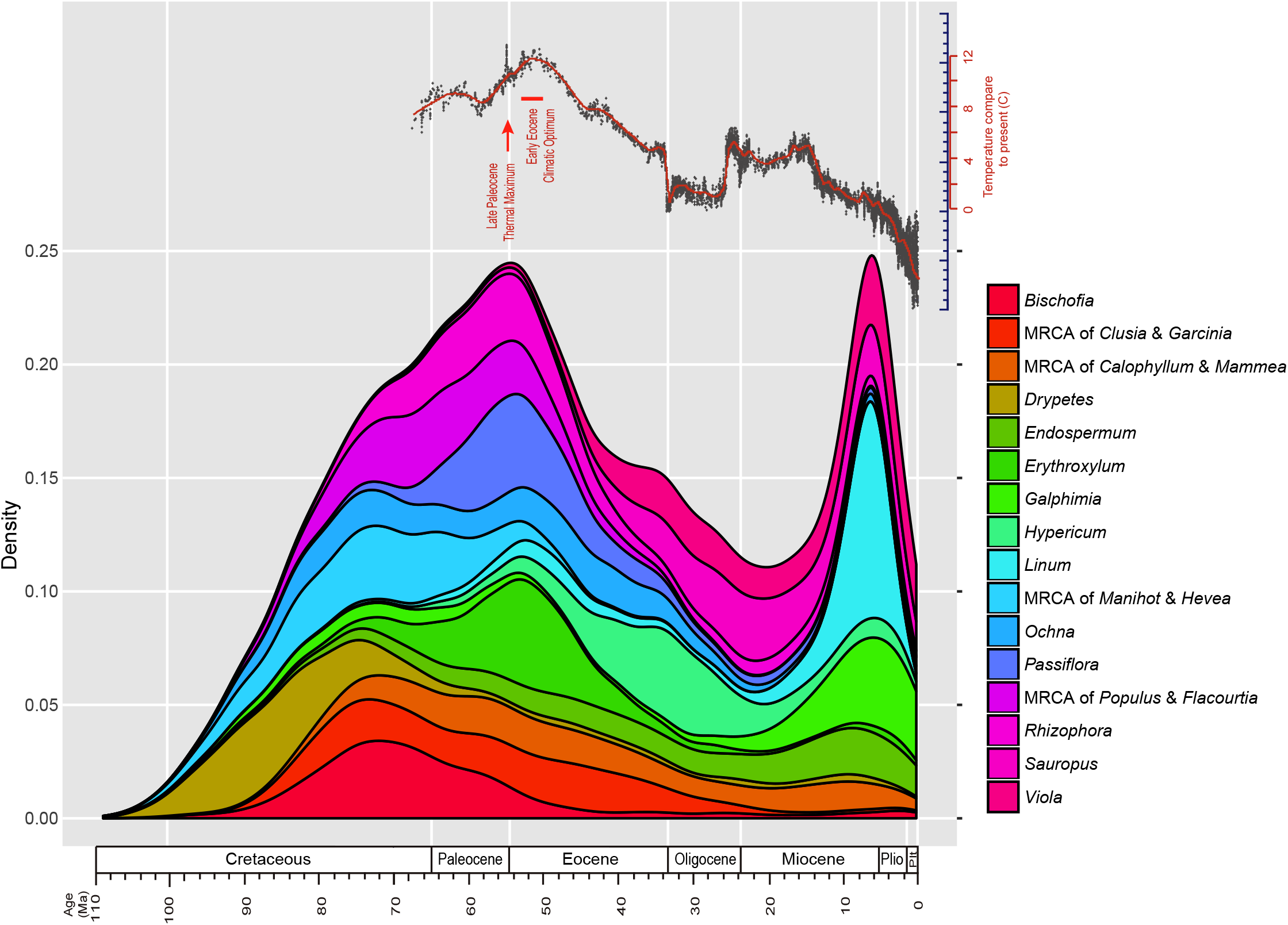
Age distributions of WGDs among clades of Malpighiales. Density of divergence time of duplicated genes by taxa are plotted in millions of years (Ma). Varying colors refer to different clades exhibiting WGDs (*Bhesa* excluded for readability). Zachos et al. (79) curve of global temperature fluctuations during the Cenozoic is redrawn at the top.

These findings are intriguing because previous studies have identified WGDs associated with major global and climatic upheavals, especially around the earlier KT boundary (~65 Ma) when a large meteor impacted off the Yucatán Peninsula disrupting the global climate, causing a major reorganization of the terrestrial biota (16). Similarly, the late Miocene-early Pliocene (~10-5 Ma) has been implicated as another period of climatic instability when WGDs were ^pervasive. The expansion of C^4 ^grassland as a result of widespread global^ aridification (59, 60) in particular, has been inferred to be correlated with numerous polyploidizations in grasses, which are among the most important members of these arid and cooler habitats that bear their namesake, i.e., grassland and steppe biomes (18). In contrast, relatively little is known about WGDs during the Eocene. This is surprising because the Eocene-Paleocene transition is associated with an extended and prolonged period of intense warming. Most notably, the Palaeocene-Eocene Thermal Maximum (PETM; 56 Ma) and the subsequent Eocene Climatic Optimum (49 Ma) constitute the warmest and most humid period during the Cenozoic. During this time, mean global temperatures increased by 5 to 10°C due to massive release of ^13^C-depleted carbon (61, 62). This dramatic climate upheaval is thought to have stimulated profound reshuffling of the terrestrial biome spurring plant migrations, extensive species turnover, and accelerated species diversification in numerous plant and animal clades (63-70). Our results establish for the first time a record of at least 19 WGDs during this period, suggesting a role in adaptation during the Palaeocene-Eocene warming intervals. We hypothesize that this may be common for predominantly tropical groups, like Malpighiales (33), which are likely much more impacted by warming given the relatively tight thermal tolerances exhibited by many tropical groups (71-73).

What may have stimulated interactions that facilitated increased polyploid formation during the warmer, wetter period of the Eocene beyond the generally increased rates of angiosperm diversification during this window of time? One hypothesis, coined the ‘neutral’ scenario (74), posits that these sorts of upheavals trigger an excess of polyploids. Here, it is widely appreciated that external stimuli, temperature in particular, has a pronounced effect on unreduced gamete formation (75). In this case, it appears that both high and low temperatures can promote unreduced gametes in various taxa as demonstrated in *Arabidopsis* (76) and some roses (77), respectively. In addition, the extensive early Miocene warming and even the Miocene aridification might have significantly increased unreduced pollens, perhaps contributing to enhanced polyploid formation. This is supported by evidence of increased levels of unreduced pollens in gymnosperms and lycophytes during comparable upheavals, including during the Triassic–Jurassic (78) and Permian–Triassic (79, 80) transitions.

Although such polyploidizations might have initially arisen more neutrally, it is also possible that polyploids were adapted for survival in these changing landscapes (the ‘adaptive’ scenario) (74). Polyploids are often viewed as evolutionary dead ends owing to their small population sizes, relatively restricted distributions, high extinction risks, and seemingly sparser representation in the deep angiosperm phylogeny (24, 81, 82). However, these circumstances may not apply under less stable environments, such as during major climatic upheavals when polyploids may outperform their diploid progenitors. Here, it has been hypothesized that such genomic novelty and epigenetic repatterning may result in phenotypic variability, including variants that confer selective advantages in stressful conditions (74, 81, 83, 84). In particular, the advantages of especially allopolyploids include altered gene expression leading to hybrid vigor and increased genetic variation. Along these lines, polyploids have been reported to be more frequent at higher latitudes, higher elevations, and in xeric environments, which may mimic such upheavals (85-88). Polyploids also occur with greater frequency among invasive plants, which commonly become established on disturbed grounds (89, 90). In support of this argument, Brochmann et al. (88) reported an unexpected overabundance of recently formed polyploids in newly deglaciated areas in the Arctic. Additionally, megaflora fossils from Wyoming, United States indicate that the dynamics of plant community assembly after dramatic warming during the PETM is very similar to late and postglacial floras (63), suggesting that observations for the Arctic may represent similar responses to warmer and wetter periods during the Eocene transition. Regardless of these competing ideas, the striking propensity and clustered distribution of WGDs in time, strengthens the hypothesis that polyploidization may be an important means of lineage persistence during episodes of major global change.

*Conclusions*–These results demonstrate astonishing levels of WGDs in a spectacular radiation of tropical flowering plants. They also lend support to the model that WGDs are an important evolutionary force in vascular plants, and that such events may be especially relevant during periods of dramatic global and climate upheaval. Finally, it has not escaped our attention that these results may confound our ability to resolve recalcitrant branches in the plant tree of life. Parsing alleles among polyploid organisms to establish orthology is already a challenge to any phylogenomics workflow (91). In the face of potentially dozens of WGDs, combined with stochastic gene copy loss, however, this obstacle could become intractable depending on the nature of the phylogenetic question at hand. Devising workflows and models to accommodate such scenarios will become increasingly more relevant as we move more deeply into the phylogenomics era.

## Materials and Methods

*Taxon sampling and transcriptome sequencing–*We collected genomic and transcriptomic data for 36 species representing 21 families of Malpighiales, spanning all major clades *sensu* Wurdack and Davis (32, 45; Table S1-3). In addition, three closely related outgroups from Celastraceae (Celastrales), Elaeocarpaceae (Oxalidales), and Oxalidaceae (Oxalidales), plus four more distantly related outgroups (*Cucumis* sativus [Eurosid I], *Theobroma cacao* [Eurosid II], and *Vitis vinifera* [basal Rosid]) were used for rooting (45). We sequenced transcriptomes of 15 Malpighiales species following the protocol described by Xi et al. (92). In short, total RNA from leaf tissue was extracted using the RNAqueous and Plant RNA Isolation Aid kits (Ambion, Inc.), and treated with the TURBO DNA-free kit (Ambion, Inc.) at 37°C for four hours to remove residual DNA. The cDNA library was synthesized from total RNA following the protocols of Novaes et al. (93). Illumina paired-end libraries were prepared for cDNA following the protocols of Bentley et al. (94). Each library was sequenced in a single lane of the Genome Analyzer II (Illumina, Inc.) with paired-end 150 base pairs (bp) read lengths. We additionally included 13 annotated transcriptomes from the OneKP project (Table S2) to complete our taxon sampling for Malpighiales. These sequences were obtained following the protocol outlined by Wickett et al. (95). Finally, we also obtained whole genome sequence data from eight published genomes of Malpighiales plus three outgroup species: *Hevea brasiliensis* (Willd. ex A.Juss.) Müll.Arg., *Jatropha curcas* L., *Linum usitatissimum* L., *Manihot esculenta* Crantz, *Populus trichocarpa* Torr. & A.Gray ex Hook., *Ricinus communis* L., *Salix purpurea* L., *Salix suchowensis* W.C.Cheng ex G.H.Zhu, *Cucumis sativus* L., *Theobroma cacao* L., *Vitis vinifera* L. (Table S3).

*Transcriptome assembly–*Raw sequencing reads were first corrected for errors using Rcorrector (96). Reads marked as ‘unfixable’, generally constituting regions of low-complexity, were discarded. Sequencing and PCR adapters were identified and trimmed using TrimGlore v0.4.2 (https://www.bioinformatics.babraham.ac.uk/projects/trim_galore/). We examined the quality of trimmed reads using FastQC v0.11.5 (https://www.bioinformatics.babraham.ac.uk/projects/fastqc/) and then assembled the reads using Trinity v2.1.1 (97). We used the longest isoform from each Trinity assembly and further reduced the redundancy generated from sequencing error or alternative splicing by performing a similarity-based clustering (-c 0.99 -n 10) via CDHIT-EST v4.6.4 (98). The completeness of our assemblies was assessed by comparison against the single-copy orthologs plant database, BUSCO (99); Fig. S4). Coding regions (CDS) of each putative transcript were predicted following the transdecoder workflow (<u>https://transdecoder.github.io/)</u>. Finally, to control for transcriptome quality in our subsequent assessments of WGD, we reanalyzed our combined transcriptome and complete genome data following the methods described below with only high quality transcriptomes. Here, transcritomes with more than 40% (382/956) missing BUSCOs were removed, including *Bhesa paniculata, Flacourtia jangomas, Galearia maingayi, Ixonanthes reticulate, Podostemum ceratophyllum, Rinorea anguifera,* and *Tristellateia australasiae*.

*Gene family clustering and orthology inference–*To assign sequences into orthologous gene families, we used an integrated method that takes into account sequence similarity and species phylogeny. We first constructed whole genome/transcriptome homology scans using Proteinortho v5.13 (100) with default parameter settings. This program extends the reciprocal best blast hit method and is computationally efficient. Clusters were searched to identify gene families containing at least 22 (>60%) ingroup species. This resulted in 8465 candidate homolog clusters. This similarity-based homology search can sometimes be erroneous due to deep paralogs, misassembly, or frameshifts (101). To reduce such errors in orthology inference, we further applied a tree-based method to sort genes into orthology groups (101). This method does not rely on a known species tree, but rather iteratively searches for the subtree with the highest number of ingroup taxa to assign as orthology groups (Fig. S5). Here, we first aligned the protein sequences of each homolog cluster using MAFFT v7.299 (102) using the local alignment algorithm (--localpair --maxiterate 1000). The resulting protein alignments were converted into the corresponding codon alignments using pal2nal v14 (103). A gene family tree of each codon alignment was then reconstructed using RAxML v8.1.5 (104) with ten random starting points. To sort homologs into ortholog groups, we first pruned exceptionally long and short branches within each gene family tree because we suspected such branches to be incorrect homologs, deep paralogs, sequencing errors, and transcript isoforms. Along these lines, branches that were ten times longer than the ‘5% trimmed mean branch length’, or shorter than an absolute value of 1e-15, were pruned. The ‘5% trimmed mean branch length’ was defined as the mean branch length after discarding the lowest and highest 5% of the branch length distribution in each gene family. Orthology was then inferred based on this pruned gene family tree using the ‘RT’ method (‘prune_paralogs_RT.py’) following Yang et al. (101). The resulting 5113 orthology clusters were realigned as amino acids using the method described above. Finally, the back-translated nucleotide alignments were used for subsequent phylogenetic analysis.

*Phylogeny reconstruction and molecular dating–*To infer the phylogeny of each orthology group, we first removed sites containing >80% gaps using trimAL v1.4.15 (105). We then applied RAxML v8.1.5 to reconstruct maximum likelihood (ML) trees under the GTR+Γ model with 20 random starting points. We then filtered each gene tree to eliminate exceptionally long and short branches using the method outlined above. The remaining gene accessions were realigned and a final round of ML tree inference was conducted. Statistical confidence of each gene tree was assessed by performing 100 bootstrap (BS) replicates with branch length (-N 100 -k).

Next, we estimated molecular divergence times for each ML gene tree as well as the bootstrap trees with penalized likelihood as implemented in r8s v1.7 (106). The following four calibration points were applied to each tree: i) the root age was fixed at 109 Ma, representing the approximate age of the crown group divergence in Malpighiales (34); ii) the minimum age of stem group clusioids (including *Calophyllum macrocarpum, Clusia rosea, Podostemum ceratophyllum, Chrysobalanus icaco, Garcinia oblongifolia, Hypericum perforatum,* and *Mammea americana*) was set to be 89 Ma, representing the oldest known fossil in Malpighiales, *Palaeoclusia chevalieri* (107); iii) two additional minimum age constraints from Xi et al. (34) were used to constrain crown group euphorbioids (107 Ma) and salicoids (94 Ma). For each clade, the age constraint was placed on the most recent common ancestor of all gene accessions forming a monophyletic group for that clade. The optimal smoothing parameter for each gene tree was determined within the range of parameter space (1e-4.5, 1e4.5) by cross-validation (106). Trees were subsequently dated under the assumption of a relaxed molecular clock by applying a semi-parametric penalized likelihood approach using a truncated Newton optimization algorithm in *r8s*.

*Species tree estimation*–We inferred a single reference species tree for our analysis of WGD applying a summary coalescent method as implemented in ASTRAL v4.10.5 (108). Our method deals with gene duplication and incomplete lineage sorting. As input gene trees for our species tree analysis we utilized all 5113 gene trees derived from each orthology cluster described above. Prior to species tree inference we applied an additional branch trimming process to remove duplicated taxa from individual gene trees following Yang et al. (101). At each node where two decedent clades contain overlapping taxa, the branch with the smaller number of taxa was pruned. One hundred bootstrap replicates were conducted for our ASTRAL analyses. Molecular divergence time estimates were subsequently inferred for the species tree using the penalized likelihood method described above using a concatenated sequence matrix derived from 40 genes containing at least 34 ingroup taxa.

*Ks-based method for WGD identification*–Each species was subjected to a reciprocal BLAST search to identify putative paralogous gene pairs in their protein coding sequences. Paralogous pairs were identified as sequences that demonstrated 40% sequence similarity over at least 300 bp from a discontiguous MegaBlast search (4, 109). Each paralogous protein sequence pair was aligned using MAFFT (102) and then back translated to their coding sequences using pal2nal (103). All sites containing gaps were removed from the alignment. Ks values for each duplicate pair were calculated using the maximum likelihood method implemented in codeml of the PAML package (110) under the F3x4 model (111). To infer WGDs from the Ks distribution, we employed the one sample K-S goodness of fit test followed by 100 bootstrap resampling to assess statistical confidence (42). A significant p value (<0.05) rejected the birth-death process supporting evidence of WGD. Because peaks produced by paleopolyploidy are expected to be approximately Gaussian (3, 112), we applied the EM algorithm to fit mixtures of Gaussian distributions to our data using the normalmixEM() function in the R package mixtools (113). Estimated mean values of peak for each taxon are reported in Fig. S1. Alternative splicing can confound the signal of WGD using the Ks method. To alleviate this concern, sites containing gaps were removed from paralog alignments, thus transcript isoforms generated from alternative splicing will receive a Ks value of zero. All pairs with a Ks value of less than 0.001, which would include these transcript isoforms as well as recent tandem duplications, were discarded and not considered in the Ks distribution.

*Placing and dating WGDs using phylogenetic reconciliation and molecular divergence time estimation*–We applied a phylogenetic approach to identify more precise placements of WGDs and to determine the approximate age of these events. We first reconciled each orthology tree to the species tree under the duplication-transfer-loss model (DTL) in Notung v2.9 (46). Here, total numbers of gene duplication inferred from well-supported gene tree nodes (>70 BS) are summarized onto the species tree. For each branch in the species tree, we calculated the percentage of genes duplicated along that branch (total number of inferred gene duplications along the branch / total number of genes containing at least one descended copy on that branch [i.e., from both single and duplicated gene copies]). The range of duplicated genes along branches where WGD was inferred was 10.0% to 84.3% (Table S4). Our threshold percentage for identifying a WGD was 10.0%, which is well above the percentage identified from fully sequenced genomes that do not exhibit recent WGDs. These genomes instead show maximally only 1.7 and 2.2% of duplicated genes where tandem duplications, not WGDs, have been inferred (calculated from *Jatropha* and *Ricinus*).

Following our phylogenetic localization of WGDs we applied a customized R script to extract the divergence times from our r8s analyses to summarize the age of gene duplications along each branch of the species tree. The inferred distribution of divergence times was fitted to a mixture of Gaussian models using the R package *mixtools* as described above to estimate mean age of each WGD. In several cases, the best model inferred two WGDs along the same branch. These cases were independently supported by our phylogenetic analysis but with smaller percentages of duplications, presumably due to gene loss or missing data (Table S4). Estimated mean values of peak for each taxon are reported in Table S4 and Fig. S6. Confidence intervals for mean age of each WGD were estimated using the 100 bootstrapped trees for each gene as outlined above (Table S4).

*Gene-count based method for confirmation of WGDs*– To further assess statistical confidence for the 24 putative WGDs inferred from the Ks method and using our phylogenetic approach above, we applied a new maximum likelihood method to test the number and placement of WGDs using gene count data (43). We first tested the utility of this method by examining three independent WGDs previously identified from fully sequenced genomes using synteny analyses. These three WGDs were identified in i.) *Populus* and *Salix* (35, 47); ii.) *Hevea* and *Manihot* (37, 49); iii.) and in *Linum* (38). For this analysis, our dated species trees was first pruned to contain only species with fully sampled genomes, including these five species, plus *Jatropha* and *Ricinus* (where WGDs have not been previously detected) and *Vitis* (as outgroup). Next, a gene count matrix was summarized for all species across all ortholog trees (Fig. S7). Filtering of the gene count matrix to avoid missing data is critical for this method, which may otherwise lead to biased estimates (43). To accomplish this, we conditioned the data matrix for all gene families to contain one or more gene copies descended from the branch along which the WGD was tested. The conditional likelihoods were subsequently estimated for models with and without the WGD of interest using a prior geometric mean of 1.5 (40). After convergence of the likelihood scores for all runs, we performed a series of likelihood ratio tests to determine the significance of individual WGDs. All three previously identified WGDs were successfully identified with confidence (LRT statistic >> 9.55, probability of type I error << 0.001). In addition, *Ricinus* and *Jatropha* showed no evidence of WGD (Table S4) suggesting a low false positive rate of this method.

We tested the remaining WGDs by sequentially adding species associated with each WGD to the seven-species phylogeny. In each case, we added all of the species from which a single WGD to be tested were descended and tested for a WGD along the added branch of interest. We generated the conditioned gene count matrix as described above. The filtering process resulted in data from 721 to 4973 gene families in all tests (Table S4). In total, we found strong statistical support for 22 of the 24 hypothesized WGDs (LRT statistic > 9.55).

*Synteny based assessment of WGD in* Linum–Synteny analysis serves as the gold standard for inferring WGDs, but is only amenable to the mostly completely assembled genomes. Our phylogenetic and Ks approach identified two WGDs in *Linum*. Here, we subsampled gene families containing three or four gene copies of *Linum* consistent with two duplications. The most closely related paralogous gene pairs of *Linum* were expected to arise from the most recent WGD, while their relationship with the remaining copy(ies) arose from the more ancient WGD (Fig. S3). We then mapped these paralogs onto the genome using MCScanX_h (105) and visualized the result using RCircos (114).

*Clustering of WGD in time–*To assess whether WGDs were clustered in time, we tested for the optimum number of clusters in age distribution using the finite Gaussian mixture modeling in R package mclust (58). The EM algorithm was used for mixture estimation and the Bayesian Information Criterion (BIC) was used for comprehensive clustering. In order to assess the confidence of the clustering, we conducted the same analysis on WGD age distributions derived from 100 bootstrap replicates.

## Acknowledgments

We thank S. Edwards, G. Giribet, E. Kellogg, A. Knoll and members of the Davis laboratory for technical assistance and valuable discussions. D. Soltis provided access to some of our transcriptome data. Funding for this study came from U.S. National Science Foundation (NSF) Assembling the Tree of Life grant DEB-0622764, DEB-1120243, and DEB-0544039 (to C.C.D.) and from Harvard University. J.S.R. acknowledges financial support from the U.S. National Institutes of Health/National Institute of General Medical Sciences grant R01 GM108904. A.M.A. acknowledges financial support from Brazil National Council for Scientific and Technological Development (CNPq) PROTAX Malpighiales grant 440543/2015-0 and Research Productivity Fellowship grant 310717/2015-9.

## Supplement figure legends

**Figure S1.**
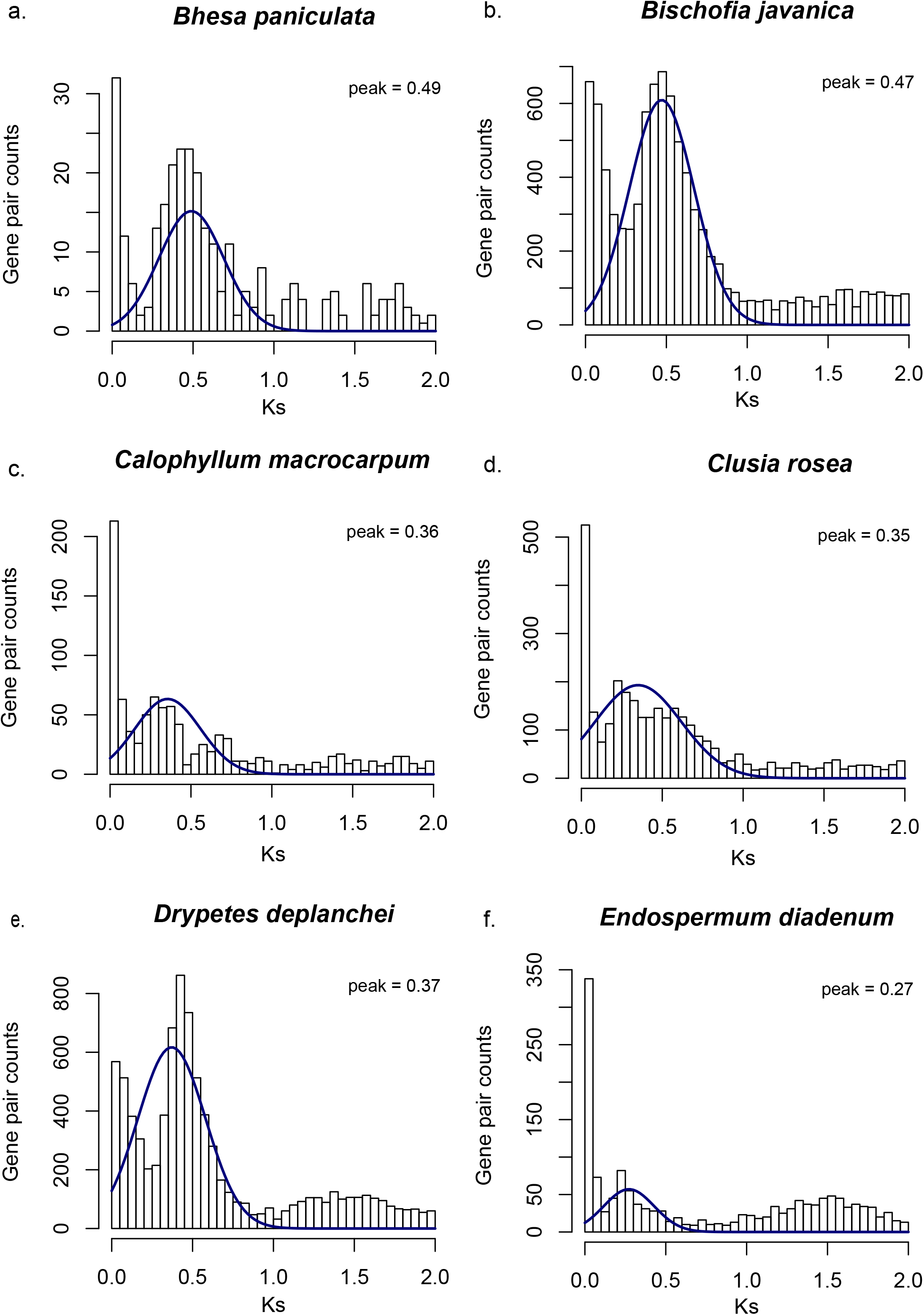
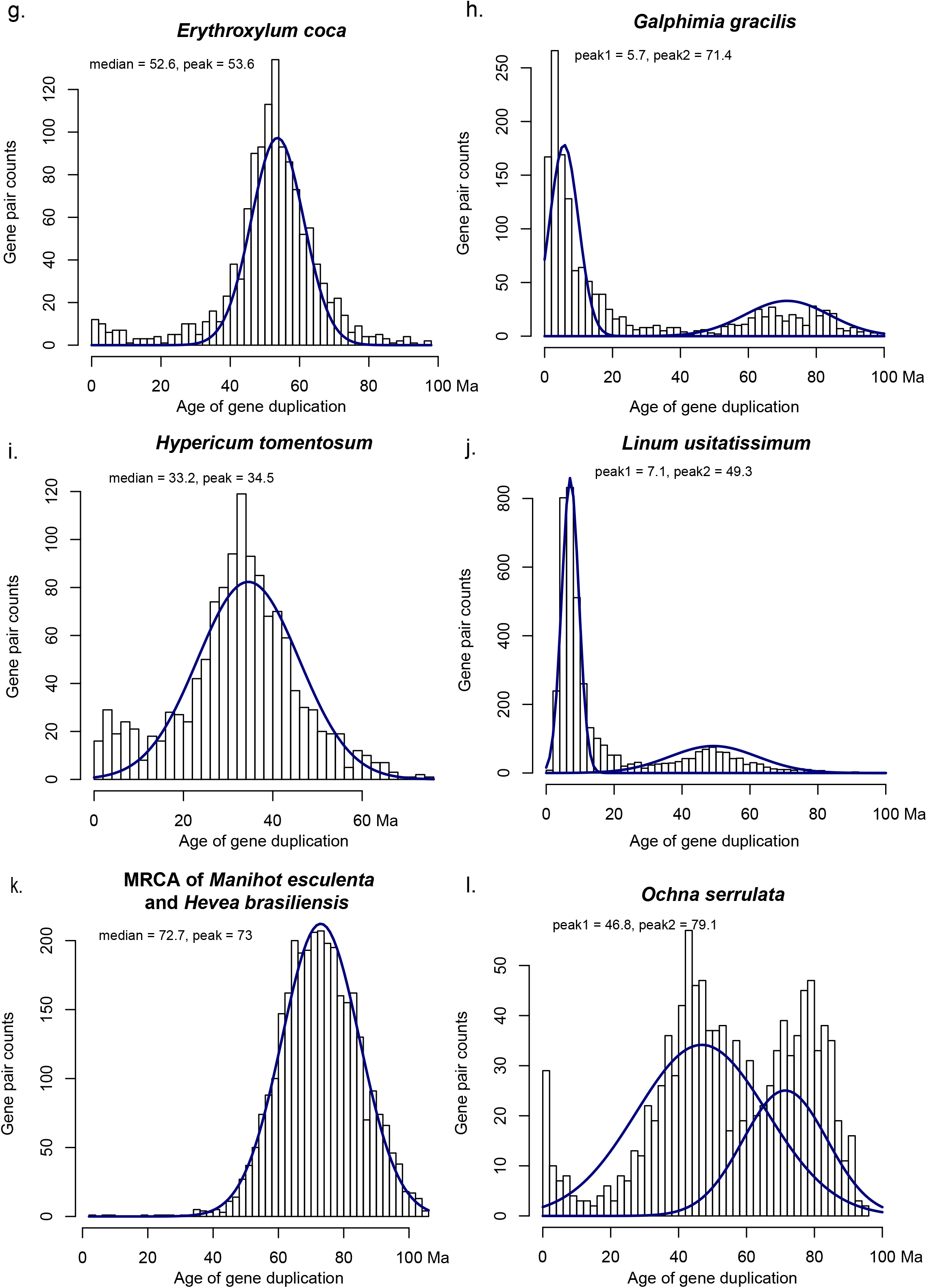
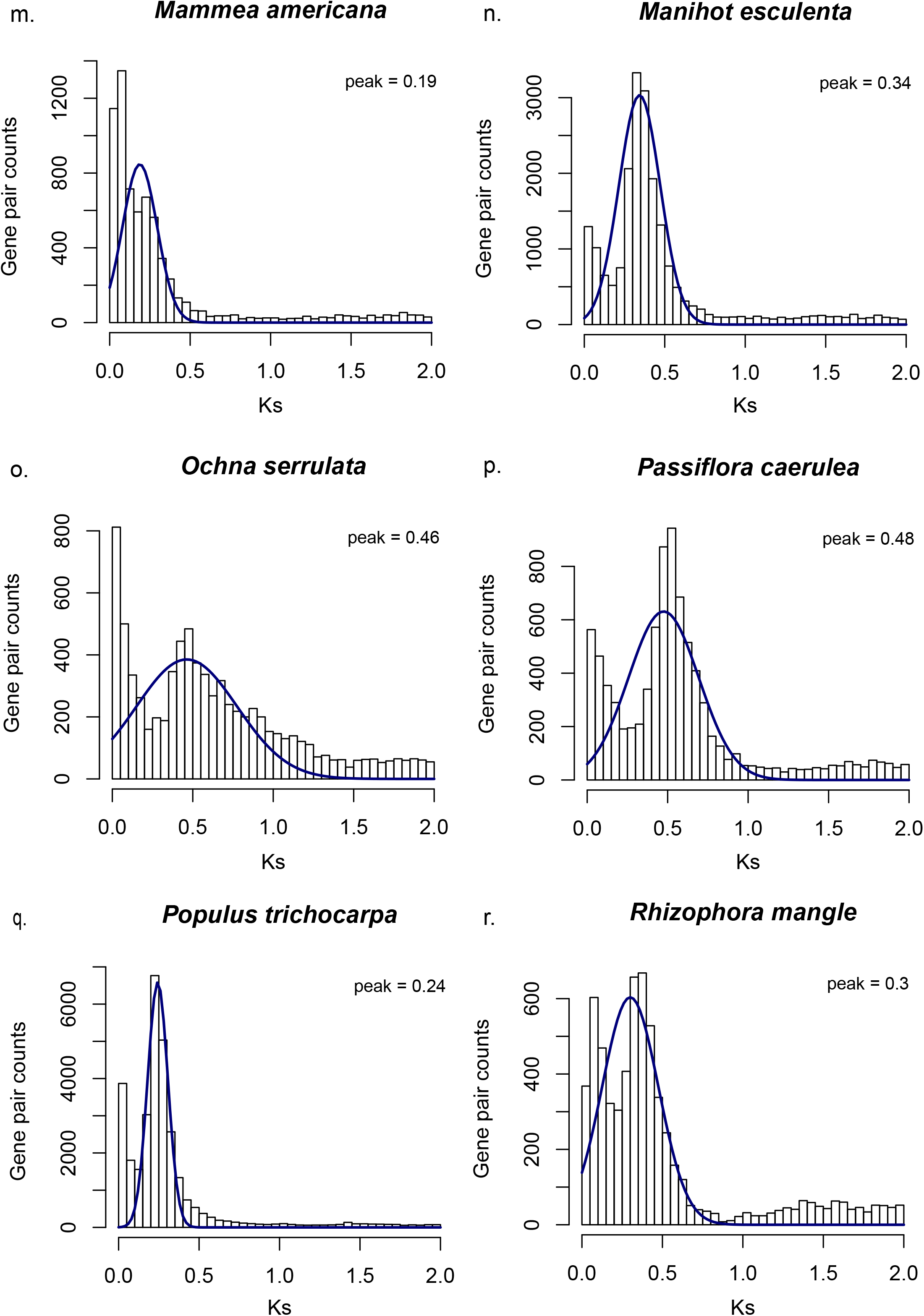
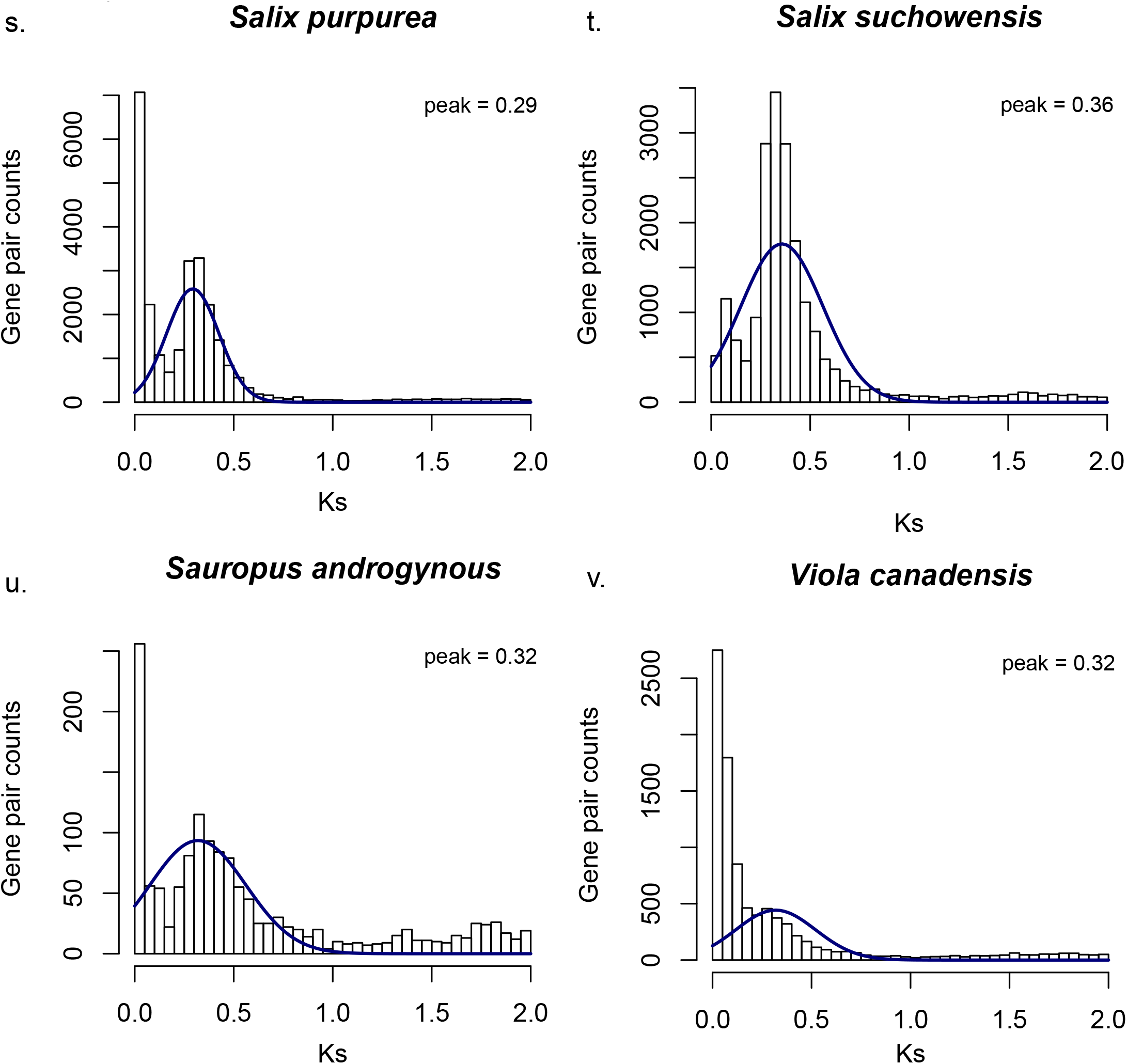
Histograms of the rate substitutions per synonymous sites (Ks) of duplicated gene pairs among 22 Malpighiales taxa showing WGDs. Taxa are as follows: a) *Bhesa paniculata* (Centroplacaceae), b). *Bishofia javanica* (Euphorbiaceae), c). *Calophyllum macrocarpum* (Calophyllaceae), d). *Clusia rosea.* (Clusiaceae), e). *Drypetes deplanchei* (Euphorbiaceae), f). *Endospermum diadenum* (Euphorbiaceae), g). *Erythroxylum coca* (Erythroxylaceae), h). *Galphimia gracilis* (Malpighiaceae), i). *Garcinia oblongifolia* (Clusiaceae), j). *Hevea brasiliensis* (Euphorbiaceae), k). *Hypericum tomentosum* (Hypericaceae), l). *Linum usitatissimum* (Linaceae), m). *Mammea americana* (Calophyllaceae), n). *Manihot esculenta* (Euphorbiaceae), o). *Ochna serrulata* (Ochnaceae), p). *Passiflora caerulea* (Passifloraceae), q). *Populus trichocarpa* (Salicaceae), r). *Rhizophora mangle* (Rhizophoraceae), s). *Salix purpurea* (Salicaceae), t). *Salix suchowensis* (Salicaceae), u). *Sauropus androgynous* (Phyllanthaceae), v). *Viola canadensis* (Violaceae). Plots j,l,n,q,s,t are derived from predicted coding sequences (CDS) using genomic data; the other plots are derived from CDS using transcriptomic data. The mean value of peak in Ks distribution is indicated on the top right.

**Figure S2.**
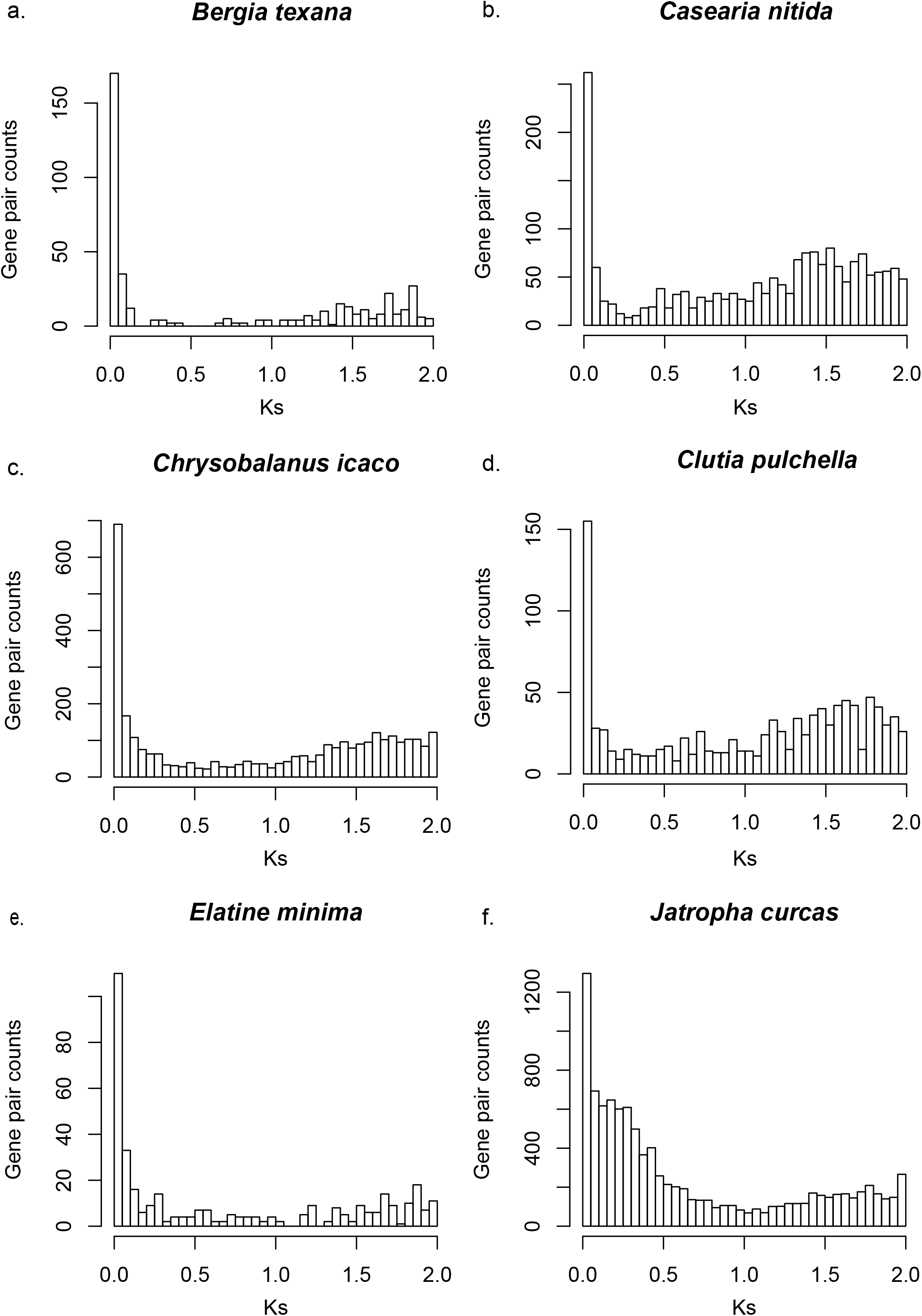
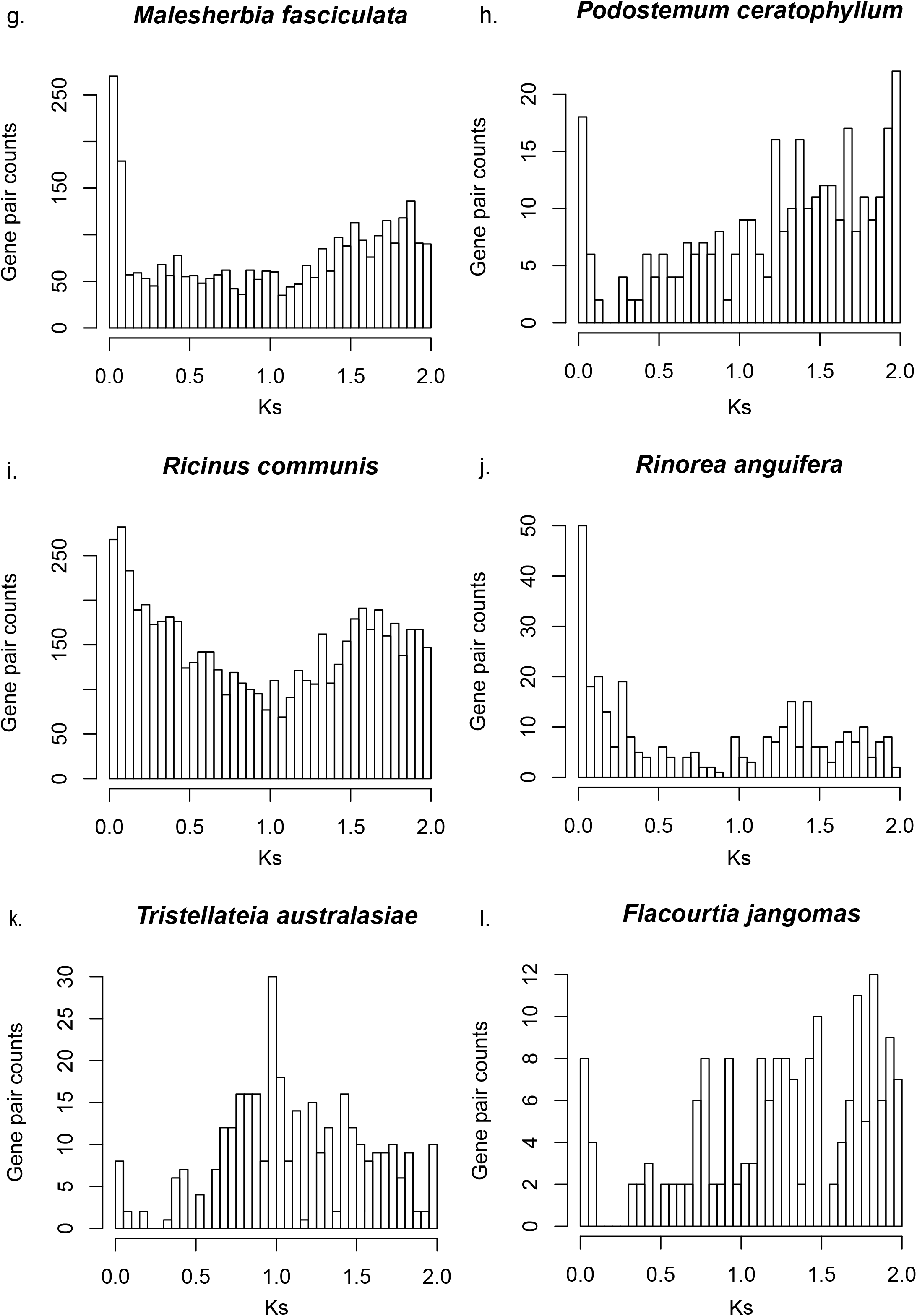
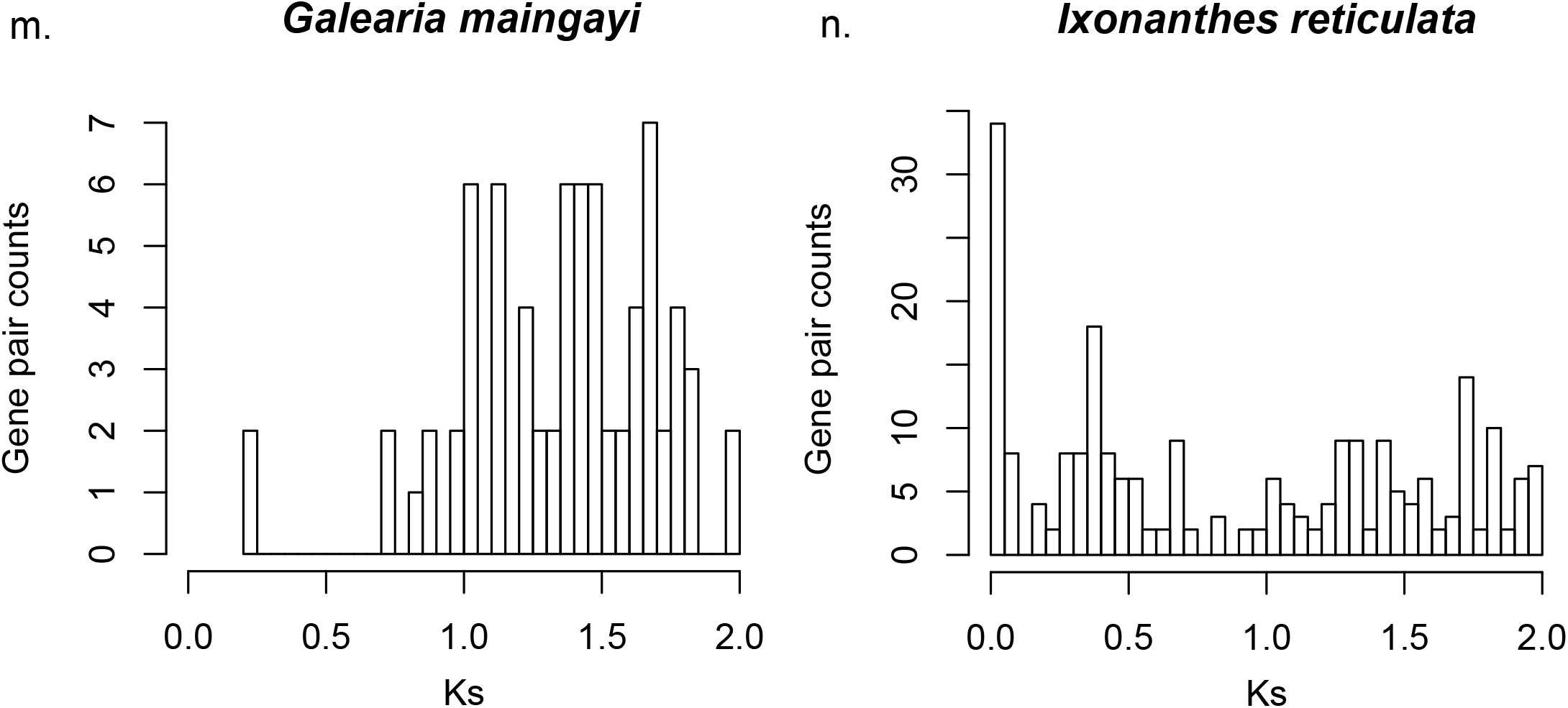
Histograms of the rate of substitutions per synonymous sites (Ks) of duplicated gene pairs among 11 Malpighiales taxa where WGD is absent (or inconclusive in the case of three taxa). Taxa are as follows: a) *Bergia texana* (Elatinaceae), b.) *Casearia nitida* (Salicaceae), c) *Chrysobalanus icaco* (Chrysobalanaceae), d) *Clutia pulchella* (Euphorbiaceae), e) *Elatine minima* (Elatinaceae), f). *Jatropha curcas* (Euphorbiaceae), g). *Malesherbia fasciculata* (Passifloraceae), h). *Podostemum ceratophyllum* (Podostemaceae), i). *Ricinus communis* (Euphorbiaceae), j). *Rinorea anguifera* (Violaceae), k). *Tristellateia australasiae* (Malpighiaceae), l). *Flacourtia jangomas* (Salicaceae), m). *Galearia maingayi* (Pandaceae), n). *Ixonanthes reticulata* (Ixonanthaceae). Plots f and i are derived from predicted coding sequences (CDS) using genomic data; the other plots are derived from CDS using transcriptomic data.

**Figure S3.**
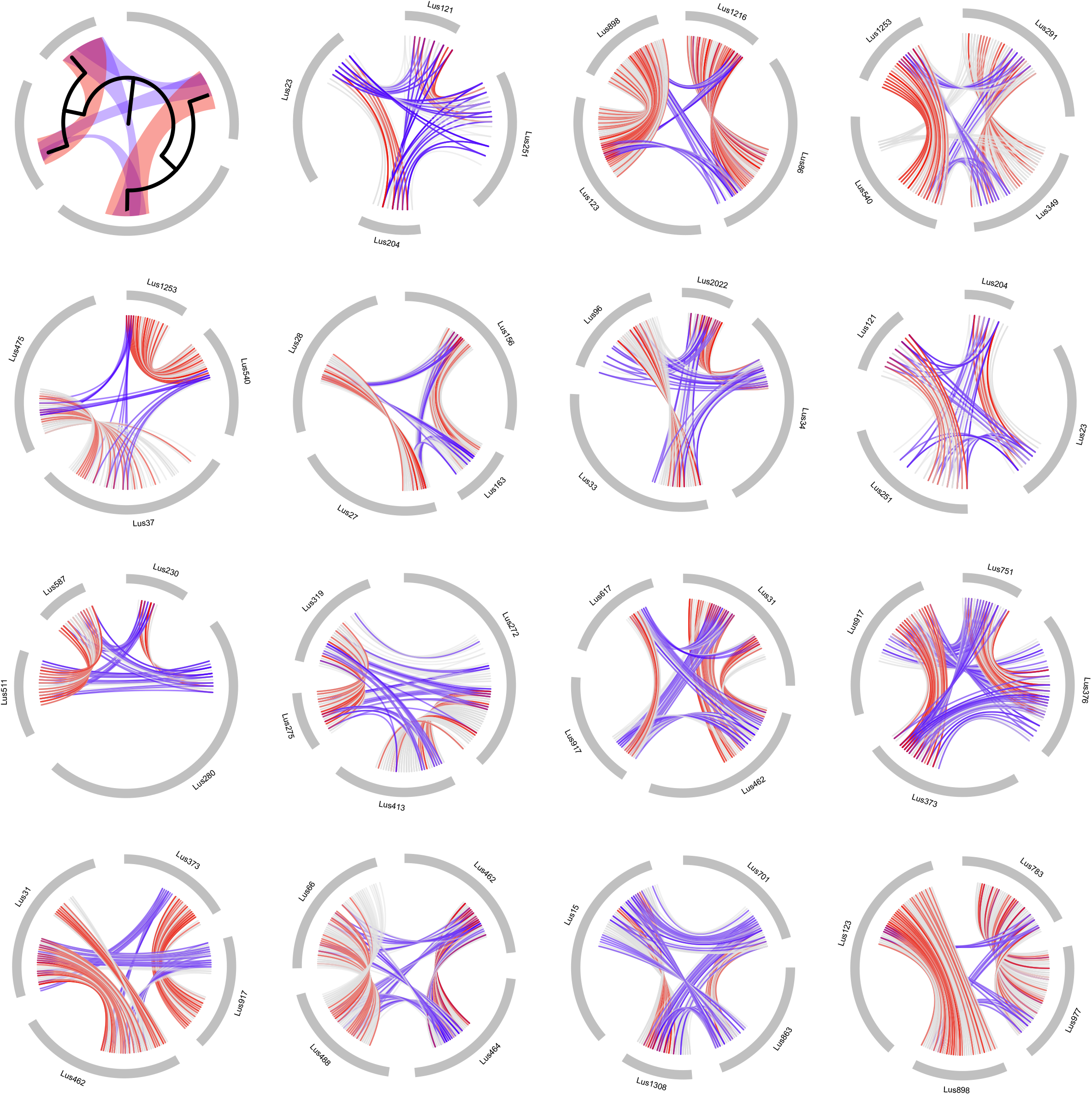
Phylogeny-guided synteny analyses demonstrate successive WGDs in *Linum.* Top left depicts a theoretical model of phylogenetic relationships for four paralogous genomic blocks resulting from two independent WGDs. Paralogous copies resulting from the most recent WGD are interconnected by red bands, while those from the older WGD are interconnected by blue bands. Fifteen large, syntenic blocks we identified (thick, grey lines) are summarized to illustrate paralogous genome regions in *Linum* that arose from two WGDs. Red and blue lines indicate paralogs of young and old WGDs, respectively, based on orthologous gene trees of the regions investigated. Thin grey lines interconnecting syntenic blocks illustrate paralogs identified by all-by-all BLAST.

**Figure S4.**
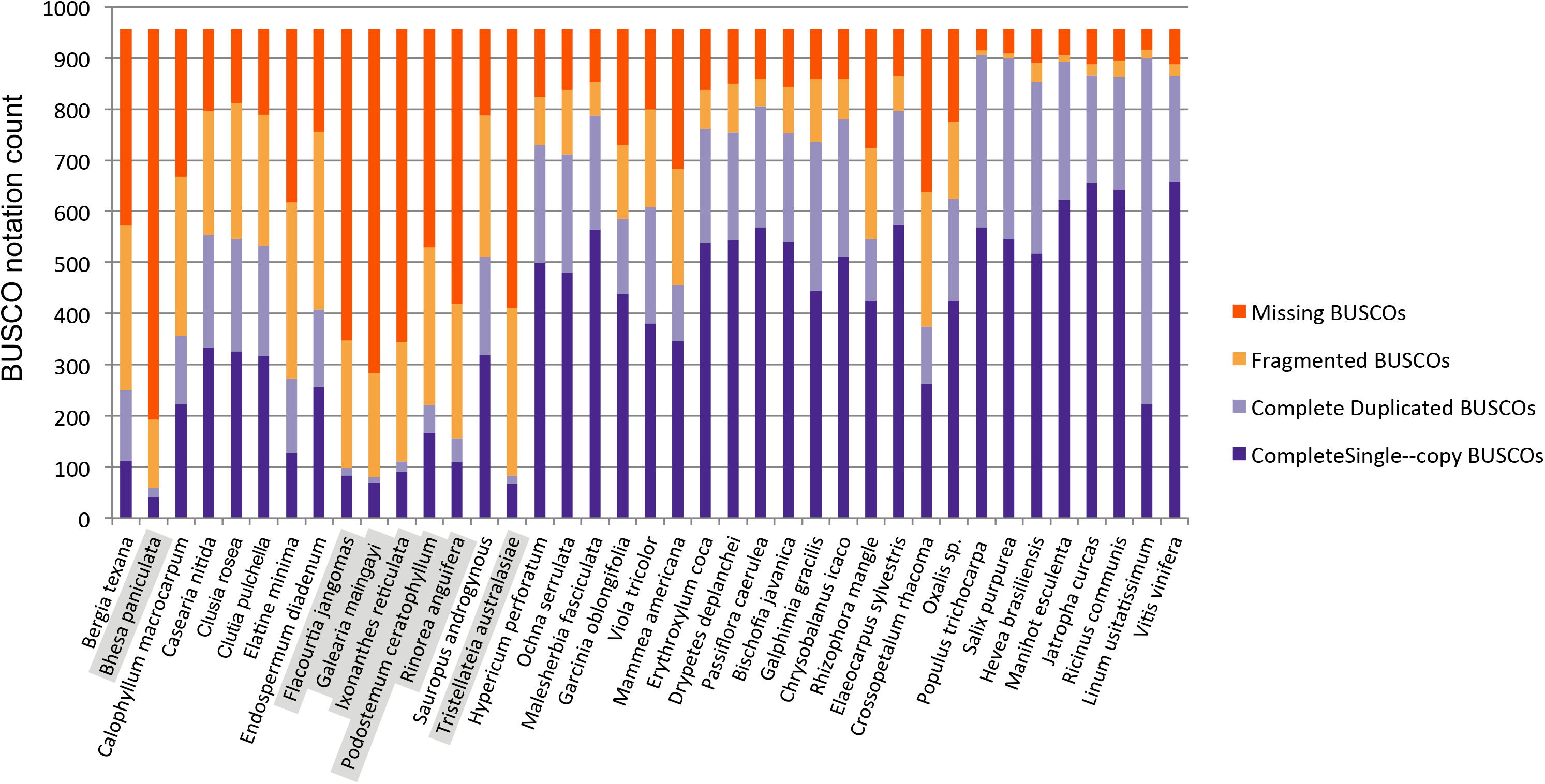
BUSCO assessment of transcriptome and genomic completeness of 42 sampled species. *De novo* assemblies and OneKP data sets are compared to the near-universal single-copy orthologs (BUSCOs). The first 15 columns represent newly generated transcriptomes for this study (from left, including *Bergia texana* to *Tristellateia australasiae*). The subsquent 16 columns represent transcriptomes acquired from the OneKP project (*Hypericum tomentosum* to *Oxalis sp.*). The last eight columns represent complete genomes for comparison (*Populus trichocarpa* to *Vitis vinifera*). Seven species with names in grey shade indicate species for which more than 40% of BUSCOS were missing. These were taxa removed in our more conservative analyses of WGD containing only high quality transcritomes and genomes (see Materials and Methods).

**Figure S5.**
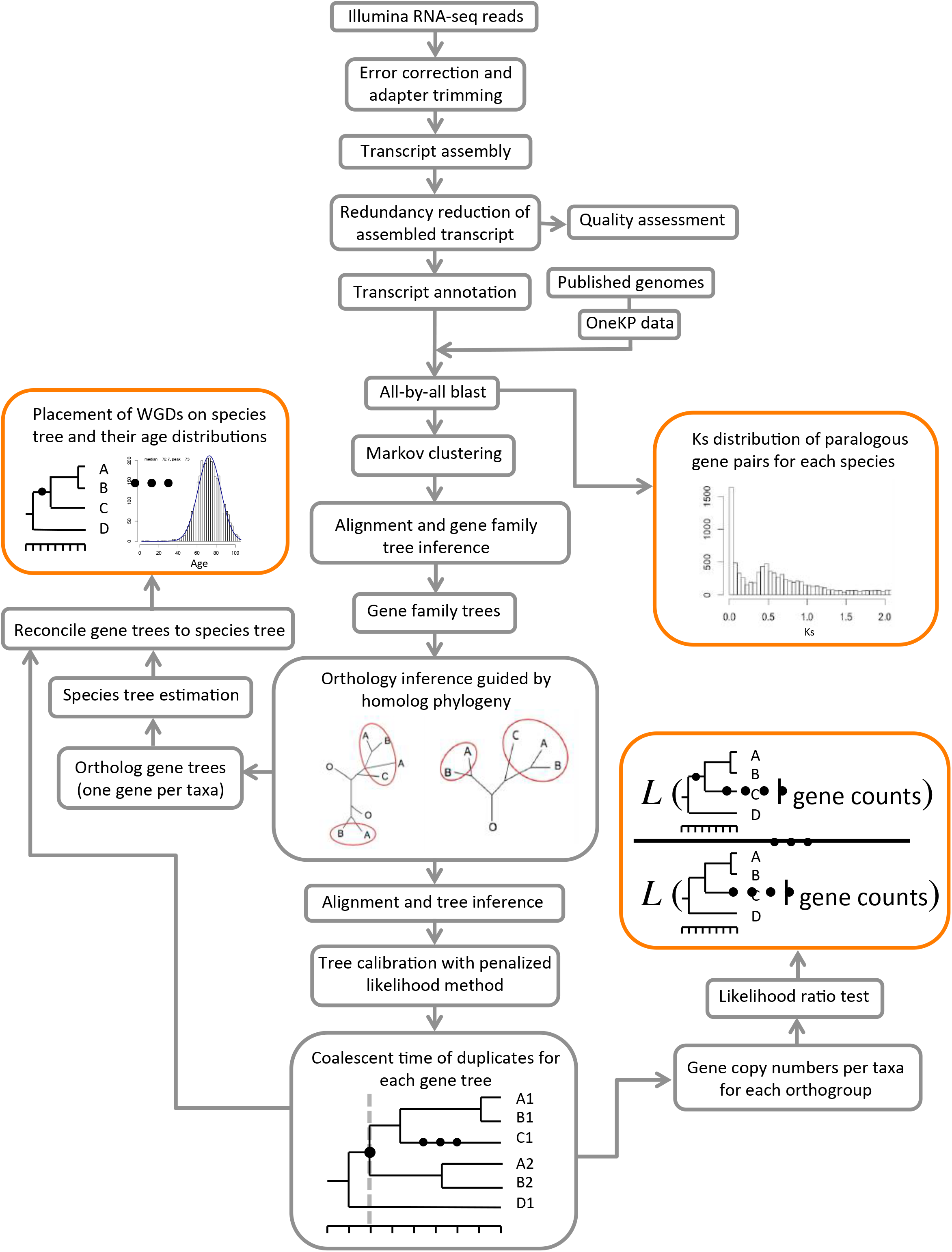
Bioinfomatic pipeline depicting transcriptome assembly, homology and orthology inferences, and three methods for WGD identification, placement, and dating.

**Figure S6.**
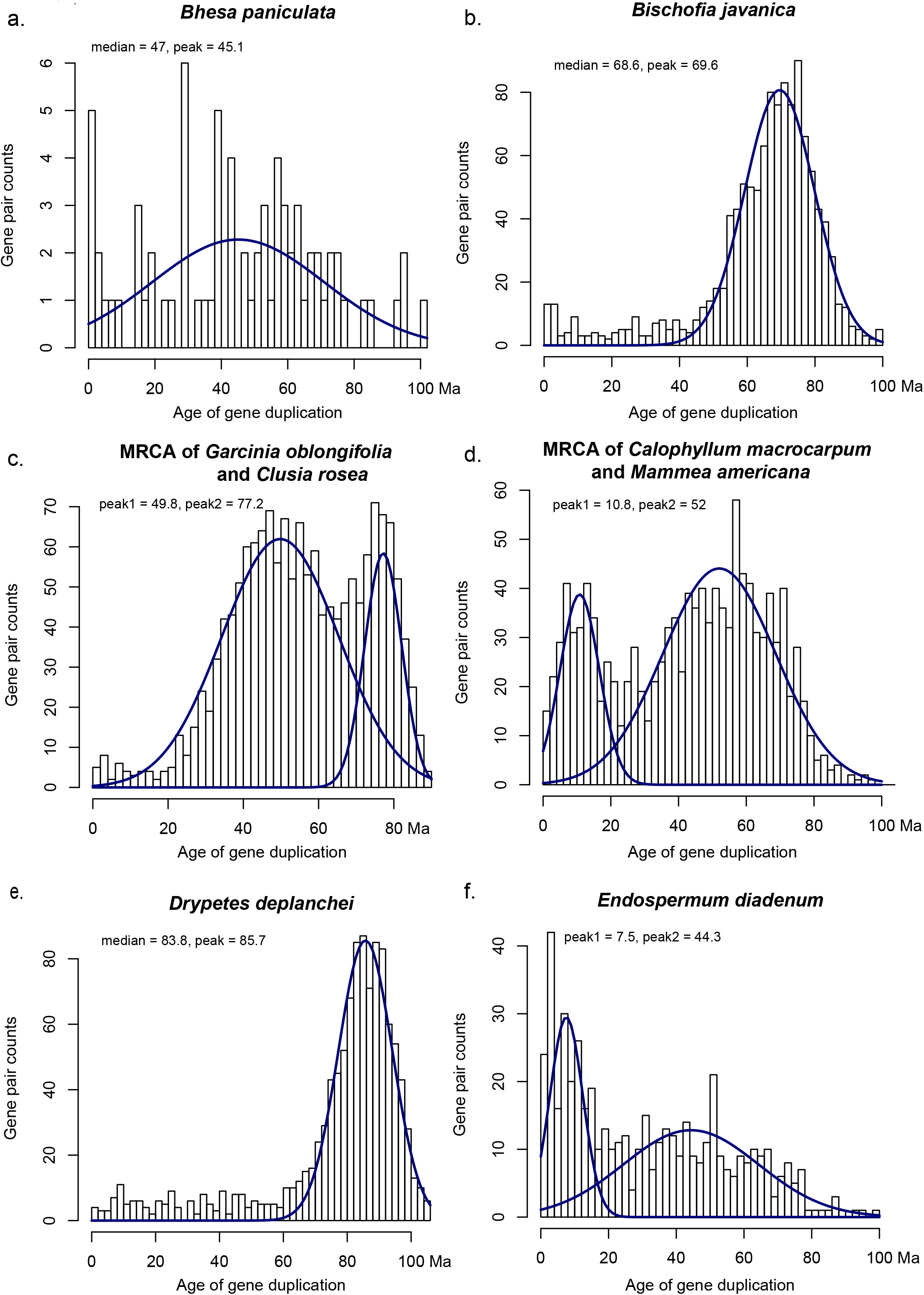
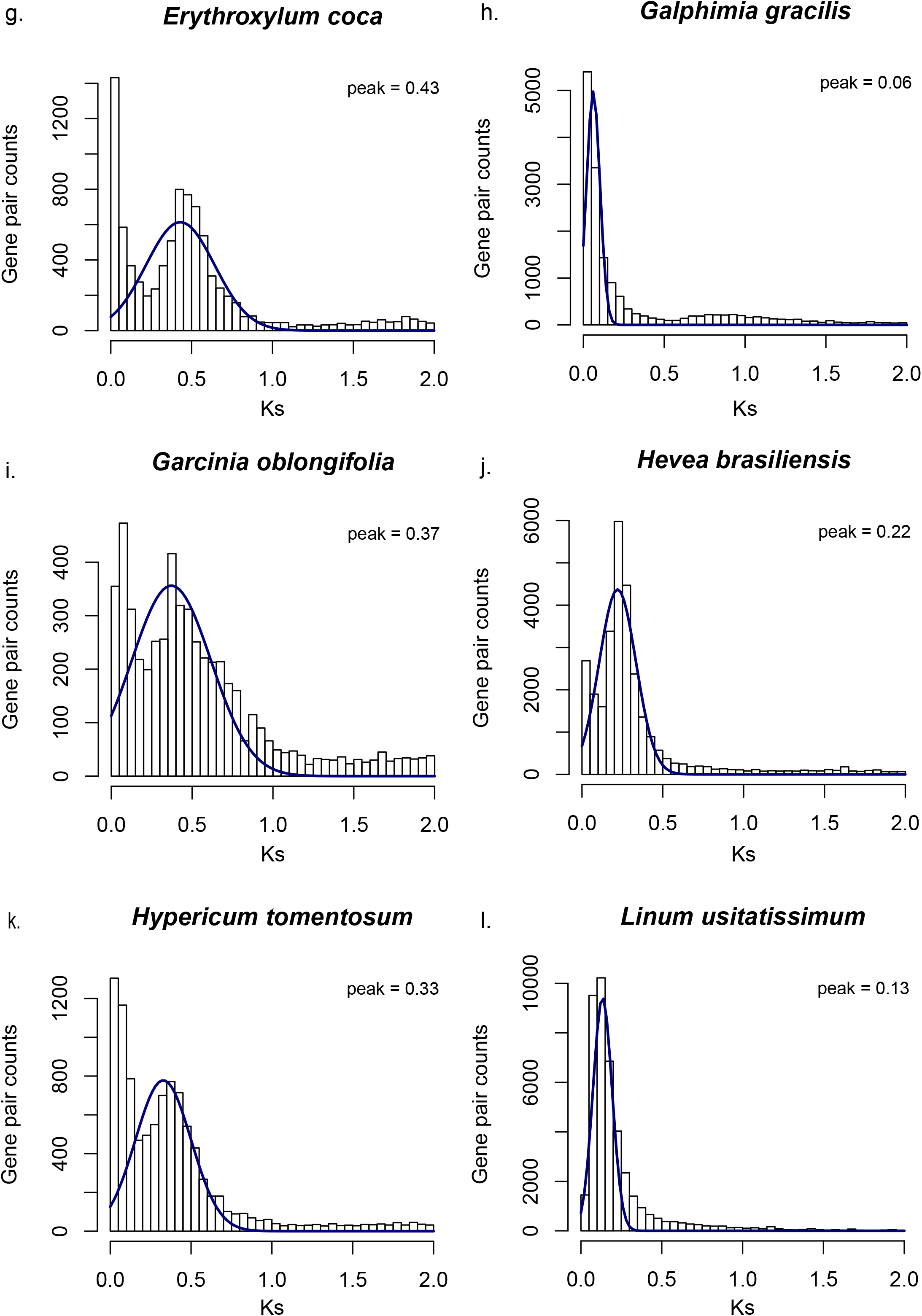
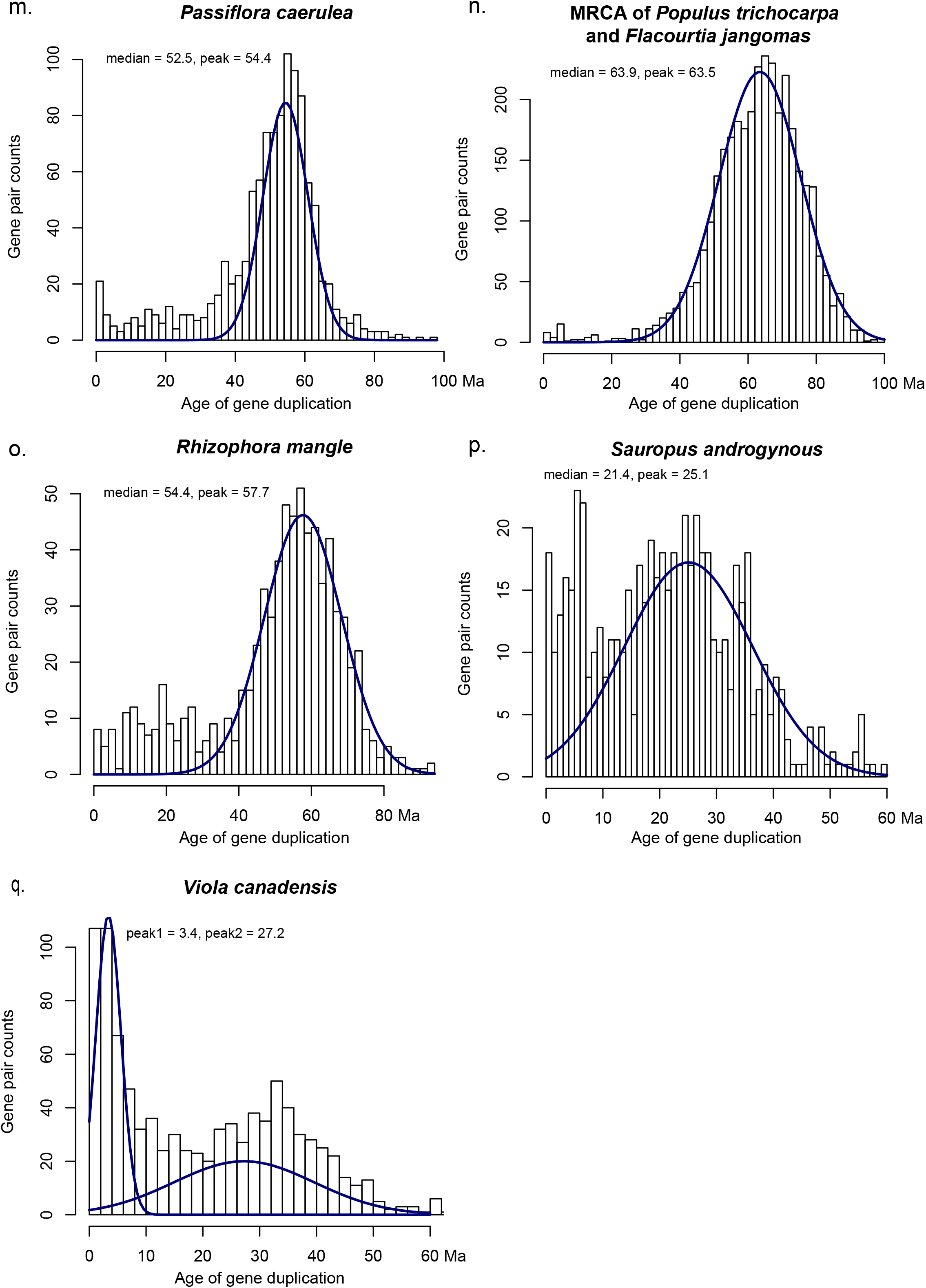
Histograms depicting divergence time estimations inferred using penalized likelihood for 22 species that exhibited WGDs (summarized in Fig. S1). Distributions are fitted to a Gaussian mixture model. Mean value(s) of mixture model(s) is reported in the top left corner of each histogram plot.

**Figure S7.**
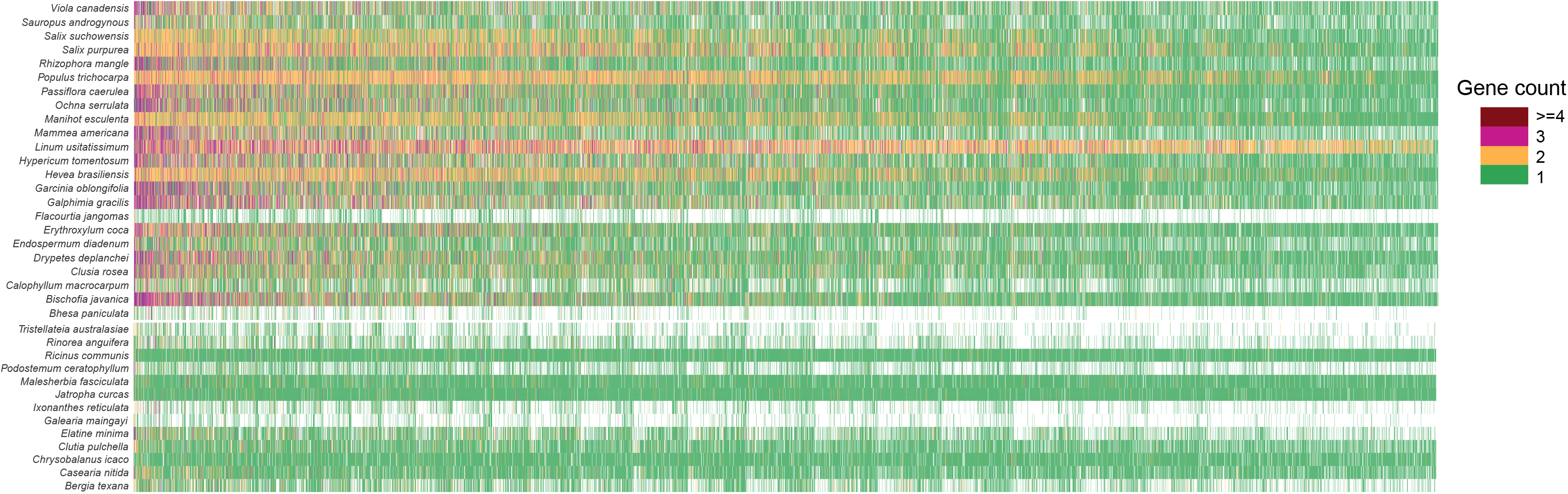
Gene count matrix for 5113 ortholog groups of 36 Malpighiales species. The first 23 species (from top to bottom) are inferred to result from WGD events using our phylogenomic reconciliation (but see Results and Discussion for *Flacourtia*); WGD is inferred to be absent in the remaining 13 species. Green, orange, purple, and brown cells represent orthogroups with one, two, three, and four gene copies for the species, respectively. Blank cells represent missing data.

**Table S1.** Voucher and GenBank information for 15 species in Malpighiales used for *de novo* transcriptome assembly. Clade name identifier *sensu* Xi et al. (2012).

| Clade name | Family | Species | Voucher | Collection Locality | GenBank Accession Number |
| --- | --- | --- | --- | --- | --- |
| Malpighiales - malpighioids | Elatinaceae | *Bergia texana* Seub. ex Walp. | Zhang, Ahart, Bartholomew 84 (A) | Butte County, California, US | ##### |
| Malpighiales - malpighioids | Centroplacaceae | *Bhesa paniculata* Arn. | Zhang and Boufford 160 (A) | Negeri Sembilan, Malaysia | ##### |
| Malpighiales - clusioids | Calophyllaceae | *Calophyllum macrocarpum* Hook.f. | SM303 | Rimba Ilmu Botanic Garden, Malaysia | ##### |
| Malpighiales - salicoids | Salicaceae | *Casearia nitida* (L.) Jacq. | FTG 72496 | Rimba Ilmu Botanic Garden, Malaysia | ##### |
| Malpighiales - clusioids | Clusiaceae | *Clusia rosea* Jacq. | Chase 341 (NCU) | Rimba Ilmu Botanic Garden, Malaysia | ##### |
| Malpighiales - euphorbioids | Peraceae | *Clutia pulchella* L. | Chase 5876 (K) | Rimba Ilmu Botanic Garden, Malaysia | ##### |
| Malpighiales - malpighioids | Elatinaceae | *Elatine minima* (Nutt.) Fisch. & C.A.Mey. | Voss 11739 (MICH) | Gemini lake, North Michigan, US | ##### |
| Malpighiales - euphorbioids | Euphorbiaceae | *Endospermum diadenum* (Miq.) Airy Shaw | SM338 | Rimba Ilmu Botanic Garden, Malaysia | ##### |
| Malpighiales - salicoids | Salicaceae | *Flacourtia jangomas* (Lour.) Raeusch. | SM289 | Rimba Ilmu Botanic Garden, Malaysia | ##### |
| Malpighiales - pandoids | Pandaceae | *Galearia maingayi* Hook. f. | SM315 | Rimba Ilmu Botanic Garden, Malaysia | ##### |
| Malpighiales - linoids | Ixonanthaceae | *Ixonanthes reticulata* Jack | SM326 | Rimba Ilmu Botanic Garden, Malaysia | ##### |
| Malpighiales - clusioids | Podostemaceae | *Podostemum ceratophyllum* Michx. | Horn & Wurdack s.n. (DUKE) | Rimba Ilmu Botanic Garden, Malaysia | ##### |
| Malpighiales - parietal clade | Violaceae | *Rinorea anguifera* Kuntze | SM307 | Rimba Ilmu Botanic Garden, Malaysia | ##### |
| Malpighiales - phyllanthoids | Phyllanthaceae | *Sauropus androgynus* Merr. | SM298 | Rimba Ilmu Botanic Garden, Malaysia | ##### |
| Malpighiales - malpighioids | Malpighiaceae | *Tristellateia australasiae* A.Rich. | Zhang 163 (A) | Cult. OEB, Harvard U. | ##### |

**Table S2.** Taxa sampled from the OneKP data set and the corresponding OneKP library ID.

| Clade or grade | Family | Species | OneKP library ID |
| --- | --- | --- | --- |
| Malpighiales - phyllanthoids | Phyllanthaceae | *Bischofia javanica* Blume | VNMY |
| Malpighiales - clusioids | Chrysobalanaceae | *Chrysobalanus icaco* L. | ZBVT |
| Malpighiales - putranoids | Putranjivaceae | *Drypetes deplanchei* (Brongn. & Gris) Merr. | RVGH |
| Malpighiales - rhizophoroids | Erythroxylaceae | *Erythroxylum coca* Lam. | RPPC |
| Malpighiales - malpighioids | Malpighiaceae | *Galphimia gracilis* Bartl. | XPBC |
| Malpighiales - clusioids | Clusiaceae | *Garcinia oblongifolia* Champ. ex Benth. | FWCQ |
| Malpighiales - clusioids | Hypericaceae | *Hypericum perforatum* L. | BNDE |
| Malpighiales - salicoids | Malesherbiaceae | *Malesherbia fasciculata* D.Don | COAQ |
| Malpighiales - clusioids | Calophyllaceae | *Mammea americana* L. | NFXV |
| Malpighiales - ochnioids | Ochnaceae | *Ochna serrulata* (Hochst.) Walp. | CKDK |
| Malpighiales - salicoids | Passifloraceae | *Passiflora caerulea* L. | SIZE |
| Malpighiales - rhizophoroids | Rhizophoraceae | *Rhizophora mangle* L. | ZTLR |
| Malpighiales - parietal clade | Violaceae | *Viola tricolor* L. | LPGY |
| Celastrales | Celastraceae | *Crossopetalum rhacoma* Crantz | IHCQ |
| Oxalidales | Elaeocarpaceae | *Elaeocarpus sylvestris* Poir. | THHD |
| Oxalidales | Oxalidaceae | *Oxalis* sp. | JHCN |

**Table S3.**
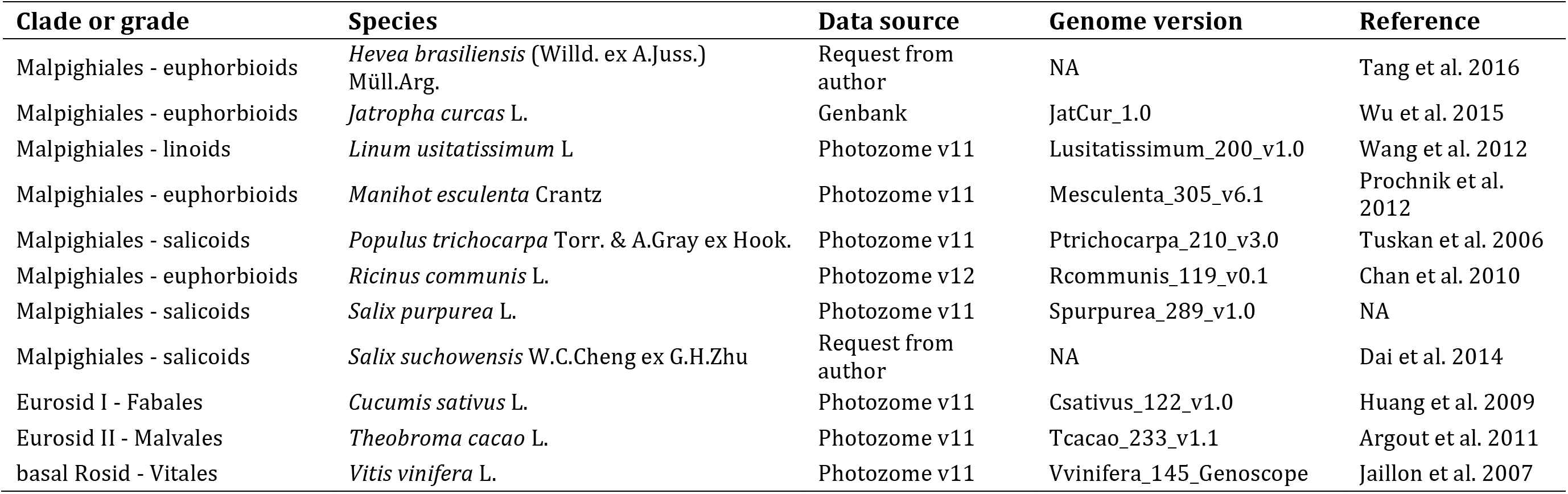
Taxa sampled with complete genomes and corresponding reference.

| Clade or grade | Species | Data source | Genome version | Reference |
| --- | --- | --- | --- | --- |
| Malpighiales - euphorbioids | *Hevea brasiliensis* (Willd. ex A.Juss.) Müll.Arg. | Request from author | NA | Tang et al. 2016 |
| Malpighiales - euphorbioids | *Jatropha curcas* L. | Genbank | JatCur_1.0 | Wu et al. 2015 |
| Malpighiales - linoids | *Linum usitatissimum* L | Photozome v11 | Lusitatissimum_200_v1.0 | Wang et al. 2012 |
| Malpighiales - euphorbioids | *Manihot esculenta* Crantz | Photozome v11 | Mesculenta_305_v6.1 | Prochnik et al. 2012 |
| Malpighiales - salicoids | *Populus trichocarpa* Torr. & A.Gray ex Hook. | Photozome v11 | Ptrichocarpa_210_v3.0 | Tuskan et al. 2006 |
| Malpighiales - euphorbioids | *Ricinus communis* L. | Photozome v12 | Rcommunis_119_v0.1 | Chan et al. 2010 |
| Malpighiales - salicoids | *Salix purpurea* L. | Photozome v11 | Spurpurea_289_v1.0 | NA |
| Malpighiales - salicoids | *Salix suchowensis* W.C.Cheng ex G.H.Zhu | Request from author | NA | Dai et al. 2014 |
| Eurosid I - Fabales | *Cucumis sativus* L. | Photozome v11 | Csativus_122_v1.0 | Huang et al. 2009 |
| Eurosid II - Malvales | *Theobroma cacao* L. | Photozome v11 | Tcacao_233_v1.1 | Argout et al. 2011 |
| basal Rosid - Vitales | *Vitis vinifera* L. | Photozome v11 | Vvinifera_145_Genoscope | Jaillon et al. 2007 |

**Table S4.** Whole genome duplications (WGDs) in Malpighiales identified with complete taxon sampling. Phylogenetic placement, percentage of gene families supporting gene duplication, log likelihood (logL, Λ>9.55 suggests significant result), and age estimations of each WGD. The last four row are species not associated with WGD based on transcriptomic (*Chrysobalanus* and *Casearia*) and genomic (*Ricinus* and *Jatropha*) data for comparison.

| WGD ID | Species | Phylogenomic reconciliation (Notung) |  | Gene-count analysis (WGDgc) |  |  | Age Estimation |  |
| --- | --- | --- | --- | --- | --- | --- | --- | --- |
| | | Raw counts (duplicated gene families/total gene families) | % | logL (duplication present) | logL (duplication absent) | $\Lambda$ | Mean (Ma) | 95% Bootstrap Confidence Interval (Ma) |
| 1 | MRCA of *Calophyllum* and *Mammea* | 510/1420 | 35.91 | -25075.79 | -25187.69 | 223.8 | 52 | 49.58636,53.31535 |
| 2 | *Mammea* | 406/3866 | 10.5 | -24594.91 | -25075.79 | 961.76 | 10.8 | 8.554801,12.279886 |
| 3 | *Hypericum* | 1229/4363 | 28.17 | -40887.4 | -41798.3 | 1821.8 | 32.5 | 33.39571,34.50537 |
| 4 | MRCA of *Clusia* and *Garcinia* | 1129/1737 | 65 | -40482.42 | -40709.61 | 454.38 | 77.2 | 57.12082,77.47120 |
| 5 |  | 361/1737 | 20.78 | -40482.42 | -40482.42 | 0 | 49.8 | 38.61796,51.11001 |
| 6 | *Ochna* | 1077/4520 | 23.83 | -42287.4 | -42962.95 | 1351.1 | 79.1 | 56.32226,79.83950 |
| 7 |  | 372/4520 | 8.23 | -42184.01 | -42287.4 | 206.78 | 46.8 | 40.02431,48.74968 |
| 8 | *Linum* | 3912/4639 | 84.33 | -37837.99 | -44349.35 | 13022.72 | 49.3 | 47.49339,59.19294 |
| 9 |  | 941/4639 | 20.28 | -37302.82 | -37837.99 | 1070.34 | 9.66 | 7.222501,7.468245 |
| 10 | *Bischofia* | 1142/4726 | 24.16 | -44497.02 | -45248.46 | 1502.88 | 68.9 | 66.3119,67.6977 |
| 11 | *Sauropus* | 594/3845 | 15.45 | -35384.61 | -35571.54 | 373.86 | 22.3 | 24.07788,32.34756 |
| 12 | *Galphimia* | 1408/4303 | 32.72 | -41050.74 | -42418.76 | 2736.04 | 56.6 | 48.00926,56.79743 |
| 13 |  | 328/4303 | 7.62 | -40916.34 | -41050.74 | 268.8 | 5.74 | 4.876005,6.030671 |
| 14 | *Rhizophora* | 725/4382 | 16.54 | -40918.22 | -41294.24 | 752.04 | 57.7 | 55.98629,60.08821 |
| 15 | *Erythroxylum* | 1199/4725 | 25.38 | -44212.69 | -44900.5 | 1375.62 | 51.5 | 53.17863,54.21989 |
| 16 | *Endospermum* | 479/3968 | 12.07 | -36473.78 | -36693.05 | 438.54 | 44.3 | 40.94213,49.50276 |
| 17 |  | 124/3968 | 3.13 | -36452.09 | -36452.09 | 0 | 7.5 | 6.076867,10.419342 |
| 18 | MRCA of *Hevea* and *Manihot* | 1528/3253 | 46.97 | -40663.02 | -43906.89 | 6487.74 | 72.8 | 73.85285,90.10148 |
| 19 | *Drypetes* | 1067/4725 | 22.58 | -44147.72 | -44819.6 | 1343.76 | 84.98 | 83.18624,85.82016 |
| 20 | *Passiflora* | 1039/4766 | 21.8 | -44590.85 | -45495.95 | 1810.2 | 49.1 | 52.43511,54.72143 |
| 21 | *Viola* | 914/4042 | 22.61 | -37482.16 | -38172.14 | 1379.96 | 27.2 | 27.31001,30.02214 |
| 22 |  | 191/4042 | 4.73 | -37414.19 | -37482.16 | 135.94 | 3.4 | 2.529189,3.495660 |
| 23 | MRCA of *Populus* and *Flacourtia* | 640/1119 | 57.19 | -34531.9 | -37494.67 | 5925.54 | 62.9 | 62.92654,63.67539 |
| 24 | *Bhesa* | 72/721 | 10 | -7024.756 | -7029.467 | 9.422 | 45.1 | 39.32973,48.67126 |
| Control | *Chrysobalanus* | 59/4738 | 1.25 | -41166.64 | -41166.64 | 0 | NA | NA |
| Control | *Casearia* | 71/4339 | 1.64 | -38567.71 | -38567.71 | 0 | NA | NA |
| Control | *Ricinus* | 81/4886 | 1.66 | -40464.76 | -37837.99 | -5253.54 | NA | NA |
| Control | *Jatropha* | 107/4973 | 2.15 | -41515.3 | -37837.99 | -7354.62 | NA | NA |

**Table S5.** Whole genome duplications (WGDs) in Malpighiales identified with more conservative taxon sampling. Phylogenetic placement, percentage of gene families supporting gene duplication, log likelihood (logL, Λ>9.55 suggests significant result), and age estimations of each WGD.

| WGD ID | Species | Phylogenomic reconciliation (Notung) |  | Gene-count analysis (WGDgc) |  |  | Age Estimation |
| --- | --- | --- | --- | --- | --- | --- | --- |
| | | Raw counts (duplicated gene families/total gene families) | % | logL (duplication present) | logL (duplication absent) | $\Lambda$ | Mean (Ma) |
| 1 | MRCA of *Calophyllum* and *Mammea* | 525/2559 | 20.52 | -40507.93 | -40873.07 | 730.28 | 48.8 |
| 2 | *Mammea* | 690/4474 | 15.42 | -40253.59 | -40507.93 | 508.68 | 12.1 |
| 3 | *Hypericum* | 1421/5467 | 25.99 | -80311.89 | -81440.86 | 2257.94 | 33.9 |
| 4 | MRCA of *Clusia* and *Garcinia* | (892+402)/2340 | 55.30 | -68371.74 | -69040.92 | 1338.36 | 76 |
| 5 |  | 402/2340 | 17.18 | -68371.55 | -68371.74 | 0.38 | 46.2 |
| 6 | *Ochna* | (781+371)/5554 | 20.74 | -81664.51 | -82387.98 | 1446.94 | 78.7 |
| 7 |  | 371/5554 | 6.68 | -81607.44 | -81664.51 | 114.14 | 47.1 |
| 8 | *Linum* | (3722+1179)/5871 | 83.48 | -79383.36 | -87342.79 | 15918.86 | 47.8 |
| 9 |  | 1179/5871 | 20.08 | -78366.2 | -79383.36 | 2034.32 | 7.3 |
| 10 | *Bischofia* | 1312/5912 | 22.19 | -87146.75 | -87949.67 | 1605.84 | 68.8 |
| 11 | *Sauropus* | 648/4348 | 14.90 | -63125.38 | -63369.66 | 488.56 | 25.6 |
| 12 | *Galphimia* | (1113+345)/5134 | 28.40 | -75243.98 | -76732 | 2976.04 | 69.9 |
| 13 |  | 345/5134 | 6.72 | -75220.94 | -75243.98 | 46.08 | 9.9 |
| 14 | *Rhizophora* | 766/5258 | 14.57 | -77393.66 | -77762.5 | 737.68 | 58.2 |
| 15 | *Erythroxylum* | 1371/5924 | 23.14 | -44212.69 | -44900.5 | 1375.62 | 53.5 |
| 16 | *Endospermum* | (547+180)/4498 | 16.16 | -65923.16 | -66287.59 | 728.86 | 47.1 |
| 17 |  | 180/4498 | 4.00 | -65834.7 | -65923.16 | 176.92 | 8.1 |
| 18 | MRCA of *Hevea* and *Manihot* | 1802/6236 | 28.90 | -84578.51 | -88456.6 | 7756.18 | 71.8 |
| 19 | *Drypetes* | 1123/5845 | 19.21 | -44147.72 | -44819.6 | 1343.76 | 84.6 |
| 20 | *Passiflora* | 1134/5956 | 19.04 | -87534.88 | -88389.97 | 1710.18 | 53.9 |
| 21 | *Viola* | (706+179)/4885 | 18.17 | -71142.56 | -71790.86 | 1296.6 | 31.9 |
| 22 |  | 179/4885 | 3.66 | -71025.62 | -71142.56 | 233.88 | 5.9 |
| 23 | MRCA of *Populus* and *Flacourtia* | 4000/6315 | 63.34 | -86816.7 | -90395.58 | 7157.76 | 60.1 |

